# Rapid Neural Reorganization during Retrieval Practice Predicts Subsequent Long-term Retention and False Memory

**DOI:** 10.1101/2020.04.02.020321

**Authors:** Liping Zhuang, Jingyi Wang, Bingsen Xiong, Cheng Bian, Lei Hao, Peter J. Bayley, Shaozheng Qin

## Abstract

Active retrieval can induce changes to the strength and content of a memory, yielding enhanced or distorted subsequent recall. But how consolidation influences these retrieval-induced seemingly contradictory outcomes remains unknown. Here we show rapid neural reorganization over eight runs of retrieval practice predicted subsequent recall. Behaviorally, retrieval practice boosted memory following a 24-hour (long-term) but not a 30-minute delay, and increased false memory at both delays. Long-term retention gains were predicted by multi-voxel representation distinctiveness in the posterior parietal cortex that increased progressively over retrieval practice. False memory was predicted by unstable representation distinctiveness in the medial temporal lobe during retrieval practice. Memory-related neural networks gradually reconfigured over retrieval practice, with the ventrolateral and medial prefrontal cortex acting as hubs for functional connections that predicted long-term retention gains and false memory outcomes respectively. Our findings demonstrate dynamic neural reorganization during retrieval practice, through which memories are arranged into discrete yet malleable representations for subsequent consolidation.

## Introduction

For centuries, humans have attempted to improve memory using mnemonic strategies. Perhaps the simplest strategy, known as retrieval practice, is through the act of retrieval, or actively recalling information as memory for that information becomes strengthened. Such retrieval practice can boost long-term retention, with robust memory gains after intervals of days or months^1^, suggesting the involvement of memory consolidation^2^. The value of retrieval practice is well recognized and widely used in education^3^, and also in the clinic to aid age-associated memory impairment^4^. Yet, each retrieval can induce transient changes in a memory’s strength and content immediately after retrieval^5^, thereby rendering the memory trace labile to be altered by current experience^6^, potentially inducing false memory^7^. However, little is known about the neurocognitive mechanisms of how memories are reorganized through retrieval practice and subsequent consolidation to produce two seemingly contradictory effects.

Psychological theories attribute the benefits of retrieval practice mainly to better access to the target memory^8^, but with different aspects of their emphasis. Specifically, a transfer appropriate processing theory emphasizes the similarity of mental operation between retrieval practice and testing phases increases the accessibility of target memory, while episodic contextual accounts highlights the similarity between the episodic contexts of practice and testing^9^. Recent neurocognitive models further suggest that retrieval practice refines neural traces and functional routes to discriminate memory representations from each other (i.e., increasing distinctiveness between memories)^10-12^. However, reactivating memories can render their neural representations labile and vulnerable to be changed by current experience^13,14^, which can initiate new learning and produce false memories^7^. Such retrieval-induced learning is believed to involve a dynamic assembly of memory-related neural circuits that reconstruct representations to meet ever-changing environmental needs^15^. Thus, practicing retrieval must actively reshape the original representations with changing connectivity among both within and between memory-related brain systems. However, there are critical gaps in our understanding of how dynamic changes in neural architectures and connectivity pathways at the time of retrieval practice can predict subsequent memory outcomes after consolidation.

One fundamental question regarding retrieval practice is how retrieval-induced transient changes in memory traces are transformed into stable representations to support subsequent long-term retention, while still being malleable enough for flexible needs. Based on systems consolidation models, memories initially rely on the medial temporal lobe (MTL, especially the hippocampus) which reinstates the cortical ensembles of encoding activity, and are then transformed into stable representations through strengthening cortical connections after days or weeks^16^. Recent evidence from humans and animal models suggests that newly acquired memories can be transformed into stable cortical engrams rapidly after encoding^17^, especially in the posterior parietal cortex (PPC)^18^. Memories for important or future-relevant information such as intensively trained ones^19^ are thought to be set as high priority^20^, by learning-induced net increases in synaptic saturation^21^ and by stronger functional connections^22^. Through subsequent offline consolidation, prioritized connections are then transformed for consolidation into long-term memory, especially during sleep^23^. By this view, retrieval practice would actively prioritize the target information for subsequent consolidation into long-term memory. A recent model, however, proposes that repeated retrieval may induce a more rapid online consolidation^10^. It is still unknown how retrieval-induced online reinstatement may interact with post-retrieval offline consolidation to affect subsequent memory.

Neuroimaging studies using localization approaches have provided useful information about the engagement of the prefrontal cortex (PFC), PPC and MTL during retrieval practice^24,25^. But they offer little insights into the multi-dimensional nature of retrieval-induced reconstruction of memory representations. Multivariate pattern analysis of fMRI data can assess active memory representations during encoding and retrieval^12,26^, using discriminative multivoxel activity patterns to characterize the distinctiveness of neural signatures within a representational space linked to individual memory events^12^. This approach allows us to assess how neural representational changes in the PFC, PPC and MTL evolve over retrieval practice and how these changes contribute to subsequent memory outcomes.

Beyond local brain regions, memory retrieval involves a complex network of widely distributed brain systems, including the PFC for retrieval search^27^, the MTL for relational memories^11,28^ and the PPC for representing specific retrieved content^29^. It is thus of interest to evaluate how the dynamic assembly of memory-related brain networks among MTL, PFC and PPC regions supports online reinstatement of memories during retrieval practice. Network neuroscience offers a graph-based network approach to provide insight into how inter-regional functional connections are gradually reconfigured over learning, including motor learning^30^, and mnemonic training^31^. To date, no study has investigated how retrieval practice reconfigures memory-related large-scale brain networks to predict subsequent memory outcome after consolidation.

Here we address the above critical questions by integrating event-related fMRI with an eight-run retrieval practice and prospective consolidation paradigm across two days (**Fig. 1A**) in combination with advanced analysis of multi-voxel representation patterns and network configurations. During the memory acquisition phase, participants were first trained to acquire 48 face-scene associations. The complex scenes associated with each face were half negative and half neutral in emotional valence for the sake of identifying a generalized effect of retrieval practice on episodic memory. Participants then underwent fMRI scanning while performing an eight-run memory practice phase with retrieval practice (RP) and no-retrieval (NR) attempts with 16 trials of each using each face as a cue. The remaining 16 trials were not presented during the memory practice phase, serving as the baseline condition. Thereafter, two cued-recall tests were given outside the scanner to assess memory retention for face-scene associations after 30-minute (short-term) and 24-hour (long-term) intervals. Memory performance of each participant was scored for their remembrance of face-scene associations (i.e., associative memory), and false memory if incorrect information were given while describing the associated scene. Multivariate and network analyses of neural activity and inter-regional connectivity involved in RP and NR conditions over 8 runs allowed us not only to track dynamic changes in memory-related neural representations and network configurations, but also to determine which specific changes predicted different memory outcomes. Based on retrieval-induced learning and systems consolidation models, we hypothesized that retrieval practice would boost long-term retention for face-scene associations while inducing false memory, and that distinct retrieval-induced changes in memory-related neural representations and network configurations would predict these two aspects of memory outcomes.

**Fig. 1.**
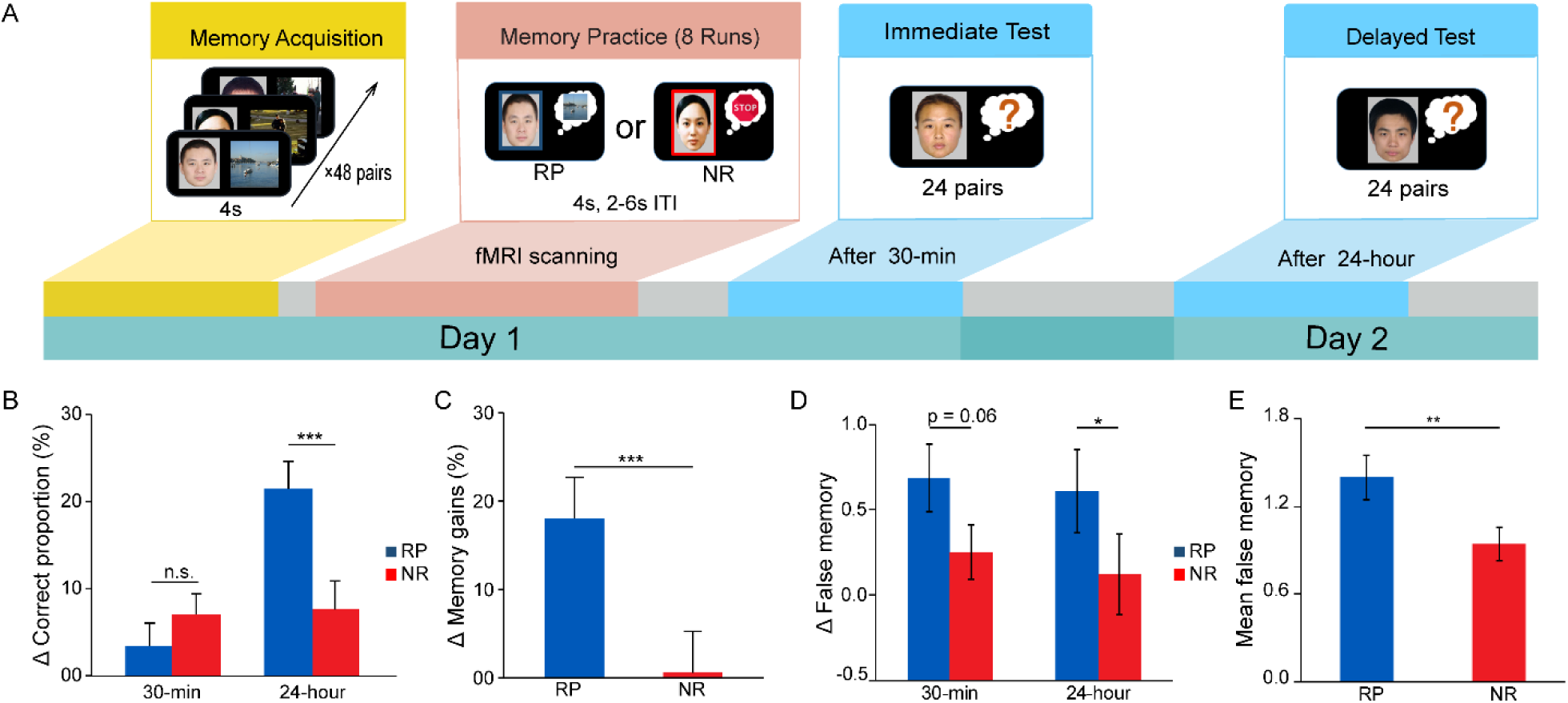
Experimental design and behavioral results. **(A)** Experimental design: The experiment consisted of three phases including memory acquisition, memory practice and subsequent memory tests. During memory acquisition, participants were trained to acquire 48 face-scene associations. During memory practice in the scanner, participants were instructed to perform either retrieval practice (RP, indicated by a blue frame) or no-retrieval (NR) attempts for the scene associated with each face over 8 runs. For each trial, participants were asked to rate the vividness of memories they recalled. Two cued-recall tests were performed to assess subsequent memory for face-scene associations after short-term (30-min) and long-term (24-hour) intervals. **(B)** Bars depict memory performance for remembering the association of each face-scene pair (i.e., associative memory) for RP and NR conditions relative to corresponding baseline. **(C)** Bars depict long-term retention gains after consolidation that is calculated with the delayed recall performance relative to the immediate recall performance. **(D)** Bars depict memory scores for false details recalled (i.e., false memory) after 30-min and 24-hour intervals for RP and NR conditions relative to corresponding baseline. **(E)** Bars depict averaged false memory scores across two days in RP and NR conditions. Notes: \**p* < 0.05; ** *p* < 0.01; *** *p* < 0.001; n.s., not significant. Error bars represent standard error of mean.

## Results

### Retrieval practice boosts long-term retention after consolidation and induces false memory

During memory practice, participants were asked to rate the vividness of memories associated with each cue for both retrieval practice (RP) and no-retrieval (NR) conditions. They reported higher vividness rating (mean ± S.D., 3.19 ± 0.43) for the RP than NR condition (1.74 ± 0.26) (t_(56)_ = 24.93, *p* < 0.001; see Supplementary **Fig. S1** for vividness ratings linked to imaging data).

We first examined the effectiveness of retrieval practice on memory retention in immediate (short-term) and delayed (long-term) tests. We observed superior long-term retention (relative to its corresponding baseline) for face-scene associations in the delayed test for RP (21.5 ± 3.1%) than NR condition (7.6 ± 3.3%) (t_(56)_= 4.37, *p* < 0.001; **Fig. 1B**). No reliable difference was observed for short-term retention in the immediate test (t_(56)_ = -1.38, *p* = 0.17; **Fig. 1B**). Moreover, we computed long-term retention gains by subtracting retention performance in the immediate test from that in the delayed test. Paired t-test revealed that long-term retention for the RP condition gained significantly higher retention scores (18.1 ± 4.6%) after consolidation than the NR condition (0.60 ± 4.6%) (t_(56)_ = 4.19, *p* < 0.001; **Fig. 1C**). In addition, we examined whether emotional valence of the associated scenes affected long-term retention gains in RP and NR conditions, by conducting 2-by-2 repeated analysis of variance (ANOVA) with Valence (Negative vs. Neutral) and Condition (RP vs. NR) as within-subject factors. This analysis revealed a main effect of Condition (F_(1,56)_ = 17.56, *p* < 0.001), but neither main effect of Valence (F_(1,56)_ = 2.68, *p* = 0.11) nor Valence-by-Condition interaction (F_(1,56)_ = 1.78, *p* = 0.19) (**Fig. S3**), indicating a similar pattern of long-term retention gains for both types of scenes associated with faces.

We then compared participants’ memory scores (relative to baseline) for false details recalled in the immediate and delayed tests for RP and NR conditions. Interestingly, we observed significantly higher false memory for the RP than NR condition in the delayed test after 24 hours (t_(49)_ = 2.49, *p* = 0.016) as well as a marginally significant effect in the immediate test after 30 minutes (t_(49)_ = 1.92, *p* = 0.06) (**Fig. 1D**). On average, false memory scores across the two delays were significantly higher in the RP (1.4 ± 0.15) than NR condition (0.94 ± 0.11) (t_(49)_ = 2.87, *p* = 0.006) (**Fig. 1E**). Together, these results indicate that retrieval practice boosts long-term retention gains for remembering face-scene associations after consolidation, and also induces false memory in general.

### Retrieval-induced changes in neural representations and their links to memory outcomes

Next, we examined retrieval-induced dynamic changes in memory-related neural representations over 8-run retrieval practices. We restricted our analyses to retrieval-related brain systems derived from a large-scale meta-analysis on the NeuroSynth platform (see Methods). To verify the validity of this mask, we conducted a set of similarity analyses for multi-voxel activity pattern during RP (or NR) condition relative to the canonical retrieval-related activation map from the NeuroSynth (**Fig. 2A**). Paired t-test revealed higher pattern similarity in the RP than NR condition (t_(49)_ = 5.51, *p* < 0.001; **Fig. 2B**). Further analyses of dynamic changes in this similarity metric over the 8 runs revealed smaller variance (variability) of neural pattern similarity in the RP than NR condition (t_(49)_ = -3.47, *p* = 0.001; **Fig. 2C**). These results indicate a greater involvement of retrieval-related brain systems in the RP than NR condition.

**Fig. 2.**
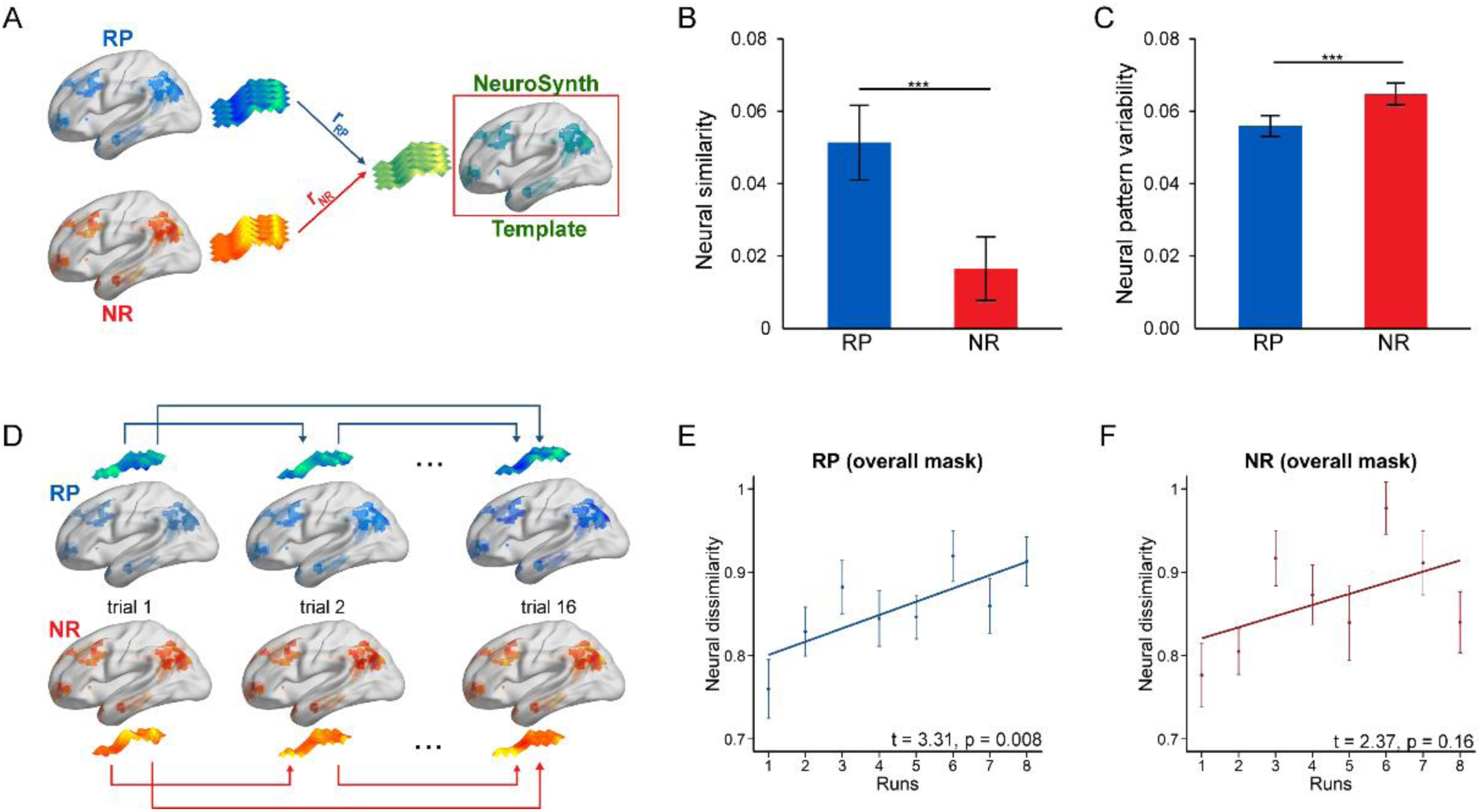
Dynamic changes in memory- related multivoxel patterns over retrieval practice. **(A)** An illustration of multivoxel pattern similarity analysis, by computing Pearson’s correlation of multivoxel neural activity pattern in RP (in blue) and NR (in red) condition with the canonical retrieval-related brain activation map derived from the Neurosynth platform (in green). **(B)** Bars depict averaged neural pattern similarity in the canonical retrieval-related brain mask across participants for RP and NR conditions separately. **(C)** Bars depict averaged variance of neural pattern similarity over 8 runs in RP and NR conditions. **(D)** An illustration of inter-trial multivoxel pattern distinctiveness, by computing pairwise pattern dissimilarity (i.e., 1 - Pearson’s correlation coefficients) among RP (or NR) trials in each run. (**E-F**) Linear regression plots show a significant linear increase in inter-trial neural distinctiveness over 8 runs in the RP but NR condition. Notes: *** *p* < 0.001. P values in (E) and (F) were adjusted with Bonferroni correction. Error bars represent standard error of mean.

We then examined dynamic changes in the distinctiveness of memory-related neural representations across individual trials over retrieval practice, by analyzing the dissimilarity of inter-trial multi-voxel neural activity patterns in the canonical retrieval-related brain mask separately for RP and NR trials in each run (**Fig. 2D**). This analysis revealed a significant linear increase in inter-trial neural pattern distinctiveness over 8 runs during the RP condition (t-test for slopes: t_(49)_ = 3.31, *p* = 0.008 after Bonferroni correction for multiple comparisons, referred as “corrected” hereafter) (**Fig. 2E**), but not during the NR condition (t_(49)_ = 2.37, *p* = 0.16 corrected) **(Fig. 2F).** Additional analyses for changes in condition-specific neural pattern fidelity and trial-specific neural signatures for the RP and NR trials are provided in Supplemental Methods & Results (**Fig. S4-S5**).

Moreover, recent neurocognitive models posit that retrieval practice could refine neural traces and the functional routes to access a targeted memory representations, thus leading to memories becoming more discriminative from each other^10^. Hence, we decomposed the overall retrieval-related canonical brain mask into three systems involving the PPC, MTL and PFC (**Fig. 3A**), and computed their corresponding inter-trial neural pattern distinctiveness for RP and NR trials in each run. We observed a linear increase in neural distinctiveness over 8 runs in the RP condition only in the PPC (t-test for slopes: t_(49)_ = 3.47, *p* = 0.005 corrected), and a marginally significant effect in the PFC (t_(49)_= 2.58, *p* = 0.08 corrected), but not in the MTL (_(49)_= 1.72, *p* = 0.70 corrected) **(Fig. 3B)**. However, we did not observe any reliable linear increase for the NR condition in these three systems (all *p* > 0.16 corrected). We further observed significantly higher inter-trial neural distinctiveness in the final run than the first run in the PPC (t_(49)_ = 3.55, *p* < 0.001) and PFC (t_(49)_ = 2.91, *p* = 0.005) in the RP condition (**Fig. 3B**) but not in the NR condition (all t_(49)_ < 1.08, all *p* > 0.28). Critically, we only found the distinctiveness of the final run in the PPC positively correlated with long-term memory gains in the RP condition (r = 0.37, *p* = 0.009), but not in the NR condition (r = -0.06, *p* = 0.67) (**Fig. 3C**). Interestingly, the inter-trial neural distinctiveness of the final run in the MTL was positively correlated with false memory in the RP condition (r = 0.38, *p* = 0.01), but not in the NR condition (r = 0.13, *p* = 0.40) (**Fig. 3D**). Collectively, these results indicate that retrieval practice leads to heterogeneous dynamics of inter-trial neutral distinctiveness in the PPC, PFC and MTL, with higher distinctiveness in the PPC predictive of better long-term retention gains, and higher distinctiveness in the MTL predictive of more false memory.

**Fig. 3.**
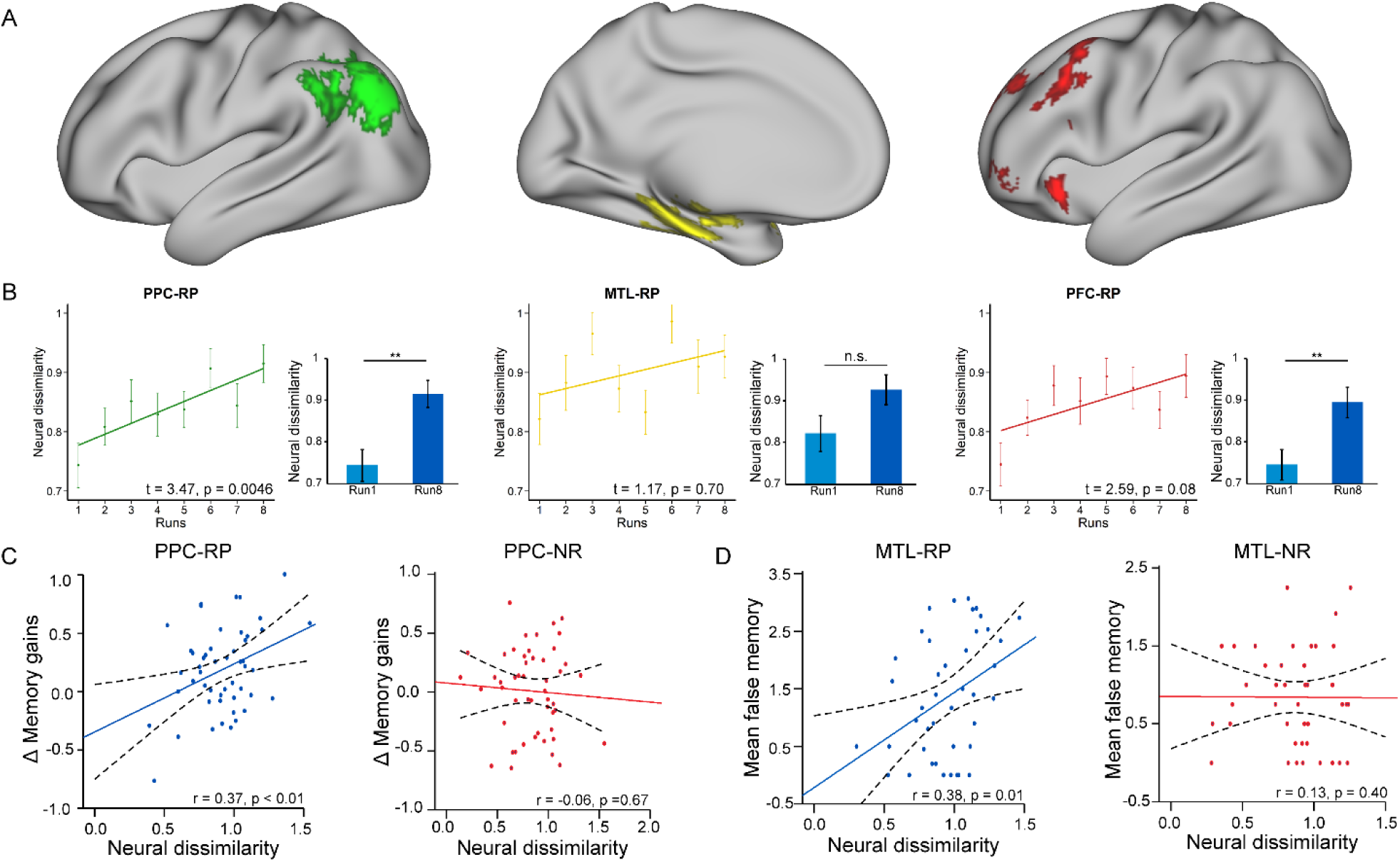
Dynamic changes in inter-trial neural pattern distinctiveness in the PPC, MTL and PFC systems and their relations to subsequent memory outcomes. **(A)** Lateral views of the posterior parietal cortex (PPC) (left panel), medial temporal lobe (MTL) (middle panel) and prefrontal cortex (PFC) (right panel) systems. (**B**) Linear regression plots show a significant linear increase in inter-trial neural pattern dissimilarity (distinctiveness) over 8 runs in the PPC, a linear trend in the PFC, and a null effect in the MTL. Bar graphs depict corresponding inter-trial neural dissimilarity between the first and the final runs in each system. **(C)** Scatter plots show positive correlation of long-term retention gains for face-scene associations with inter-trial neural dissimilarity in the final run for the RP condition (left), but not for the NR condition (right). **(D)** Scatter plots show positive correlation of false memory scores with inter-trial neural dissimilarity in the final run for the RP condition (left), but not for the NR condition (right). Notes: Beta and p values are from the least squares fitting, p values were adjusted with Bonferroni correction. Error bars represent standard error of mean.

### Retrieval-induced reorganization of brain networks to predict long-term retention gains

To track the dynamic assembly of large-scale brain networks including 15 nodes in PPC, PFC and MTL over retrieval practice, we constructed a network consisting of 15 × 15 pairwise links for each run (**Fig. 4A**), using a generalized form of context-dependent psychophysiological interactions (gPPI). Overall, we observed a significant increase in global network efficiency in the RP condition (t-test for slopes: t_(49)_= 2.78, *p* = 0.0077), but a significant decrease in the NR condition (t_(49)_= -2.05, *p* = 0.046) (**Fig. 4B**). Follow-up statistical tests revealed significantly higher slopes in the RP than NR condition (all t_(49)_ > 3.37, *p* < 0.0015) (**Fig. 4C**). Additional analyses also revealed dynamic changes in network configuration among the PPC, PFC and MTL over retrieval practice (**Fig. S7**).

**Fig. 4.**
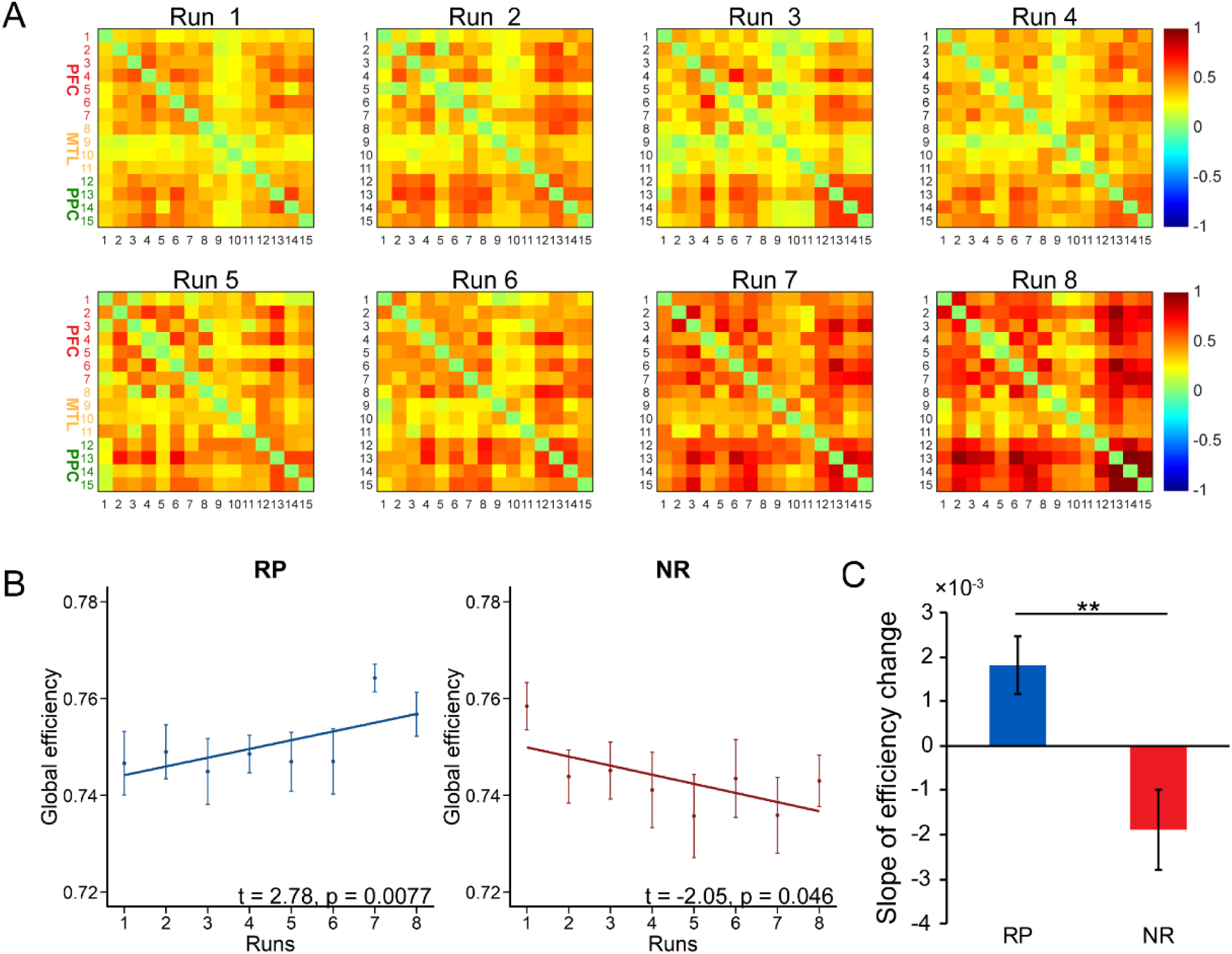
Dynamic changes in memory-related neural network configurations over retrieval practice. **(A)** Network matrices show pairwise inter-regional connectivity links over 8 runs during retrieval practice. Color bars reflect task-dependent functional connectivity strength based on a generalized form of context-dependent psychophysiological interaction (gPPI) approach. The strength of links shows a gradual increase over the course of retrieval practice. **(B)** Linear regression plots show a significant linear increase in network efficiency as a function of 8 runs for the RP condition (left), and a linear decrease over 8 runs for the NR condition (right). **(C)** Bars depict the slope of dynamic changes in network efficiency, with significantly higher in the RP than NR condition. Notes: Beta and p values are derived from the least squares fitting. ** P < 0.01. Error bars represent standard error of mean.

We then examined how dynamic reconfiguration of memory-related brain networks over retrieval practice contributed to long-term retention gains after consolidation. We implemented a network-behavior prediction analysis by training a support vector regression (SVR) model based on network features over 8 runs in the RP (or NR) condition. A feature selection procedure revealed that information derived from the top 1% links of all 8 runs as the input features provided the most robust accuracy to predict long-term retention gains (**Fig. S8A**). The selected links from each run were visualized in **Fig. 5A**.

**Fig. 5.**
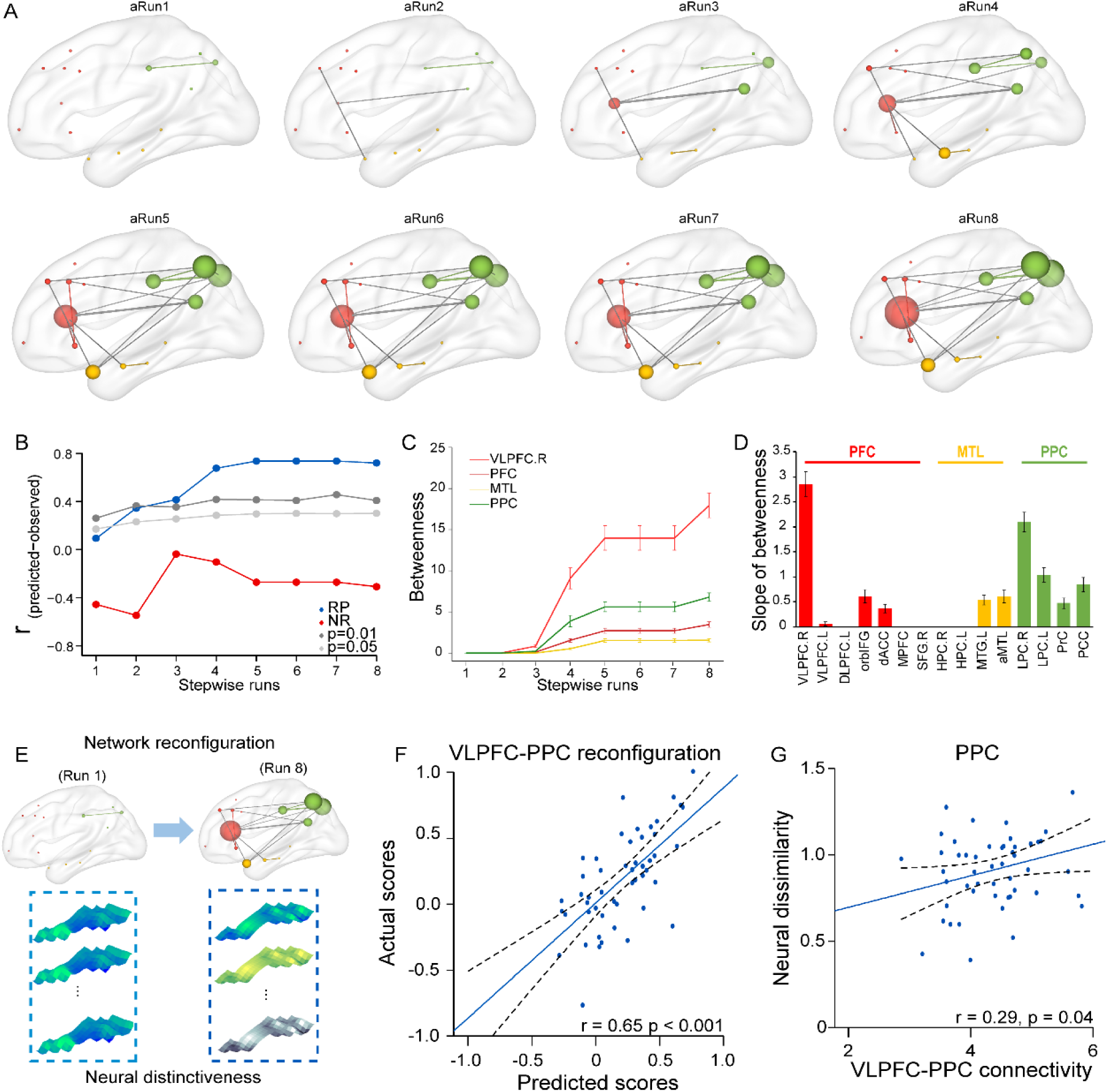
Brain network-based prediction of long-term retention gains after consolidation and network reconfigurations over retrieval practice. **(A)** Lateral views of selected links critical for predicting long-retention gains project onto a glass brain to illustrate dynamic changes in memory-related neural network configurations over 8 runs retrieval practice. Red, green and yellow nodes are regions of interests (ROIs) in the prefrontal cortex (PFC), the medial temporal lobe (MTL) and the posterior parietal cortex (PPC), respectively. **(B)** Line plots show prediction values from stepwise brain-behavior prediction analyses for long-term retention gains. Prediction values from the RP condition reach higher than 0.74 over the first 5 runs and then remain relatively stable from **runs 5** to **8** based on stepwise accumulative information as the progression of retrieval practice. The prediction values in the RP condition clearly outperforms the NR condition as well as the randomly-permutated significance at *p* < 0.01. **(C)** Line plots show betweenness centrality over 8 runs for the RP condition in the ventrolateral prefrontal cortex (VLPFC), MTL, PFC and PPC systems. (**D**) Bars depict the slopes of dynamic changes in betweenness centrality over 8 runs in 15 nodes for the RP condition. Notes: Abbreviations for 15 nodes are listed in Supplemental Texts. Error bars represent standard error of mean. **(E)** An illustration of the VLPFC- and PPC-dominated brain network reconfigurations and fine-tuned inter-trial neural pattern distinctiveness over retrieval practice. **(F)** Scatter plot shows a highly positive correlation of observed long-term retention gains and predicted outcomes from prediction analysis based on functional connectivity strength of the right VLPFC with PPC nodes during retrieval practice. **(G)** Scatter plot shows a positive correlation between neural pattern distinctiveness in the PPC and functional connectivity strength of the right VLPFC with PPC nodes. Notes: VLPFC, ventrolateral prefrontal cortex; PPC, posterior parietal cortex.

We then performed a stepwise prediction analysis to characterize changes in predictive values as a function of 8 runs by using selected links described above (**Fig. 5B**). The highest prediction value in the RP condition reached 0.74 (r_(predicted, observed)_ = 0.74, *p* < 0.001 permutation test) over the first 5 runs and remained at a stable level since then. This prediction in the RP condition outperformed the NR condition (**Fig. 5B**; gray lines). To illustrate the network properties in the RP condition, we performed graph theory-based network analyses using the selected links across 8 runs (**Fig. 5A**). These analyses revealed a significant increase in betweenness centrality for nodes in the PPC, PFC and MTL (t-test for slopes: t_(49)_ >17.27, all *p* < 0.001), with more prominent increase in the PPC than the PFC and MTL (both t_(49)_ > 9.6, both *p* < 0.001; **Fig. 5C**). Of the 15 nodes, the right VLPFC showed the most prominent increase far above that of the others (all t_(49)_ > 4.2, all *p* < 0.001; **Fig. 5D**). These results indicate that dynamic reconfiguration of memory-related brain networks over retrieval practice is predictive of long-term retention gains after consolidation, with the right VLPFC emerging as a hub.

### VLPFC and PPC dynamic reorganization predicts inter-trial neural pattern distinctiveness

Given that the VLPFC and PPC stood out as two critical nodes involved in dynamic reorganization of memory-related neural networks over retrieval practice, we further hypothesized that their functional coordination would facilitate the distinctiveness of neural representation patterns to predict long-term retention gains (**Fig. 5E**). To test this hypothesis, we implemented an additional prediction analysis by only using the links between the right VLPFC and PPC nodes as input features. This analysis again revealed that connectivity strength between these nodes was highly predictive of long-term retention gains (r_(predicted, observed)_ = 0.66, *p* < 0.001; **Fig. 5F**), indicating that communication ability between these regions during retrieval practice is critical for subsequent memory outcomes.

Furthermore, we examined the relationship between VLPFC network connectivity and inter-trial neural pattern distinctiveness in the PPC nodes that has been considered to be an “output buffer” for specific representations of the retrieved content(*38,42*). We observed that connectivity strength of the effective links between the VLPFC and PPC nodes was positively predictive of inter-trial neural pattern distinctiveness in the PPC in the final run (r = 0.29, *p* = 0.04; **Fig. 5G**). These results indicate that functional connectivity between the right VLPFC and PPC over retrieval practice is critical to predict inter-trial neural pattern distinctiveness in the PPC and long-term retention gains.

### MPFC and MTL dynamic reorganization during retrieval practice predicts false memory

To examine how retrieval-induced network reconfigurations contribute to false memory, we conducted network-behavior prediction analyses for false memory scores using the same procedures we used to predict long-term retention gains. Given that false memory showed a similar pattern for the 30-min and 24-hour intervals, we used the mean false memory scores across the two intervals for this prediction analysis.

A feature selection procedure again revealed that information derived from the top 1% links of all 8 runs as input features yielded the best outcome to predict false memory (**Fig. S8B**). We plotted the stepwise prediction values as a function of 8 runs, with the prediction value reaching 0.55 (r_(predicted, observed)_ = 0.55, *p* < 0.001 permutation test) across the first 5 runs and increasing to 0.74 in the final run for the RP condition. The selected features were visualized in **Fig. 6A**. This prediction in the RP condition also outperformed the NR condition (**Fig. 6B**). Unlike the right VLPFC acting as a hub to predict long-term retention gains, graph theory-based network analyses for selected links revealed a gradual increase in betweenness centrality for the PFC, MTL and PPC (t-test for slopes: t_(49)_ > 7.15, all *p* < 0.001), with the most prominent increase in the MPFC (**Fig. 6C**). The posterior cingulate cortex and the right hippocampus also showed more prominent increase in betweenness centrality than other nodes (all t_(49)_ > 3.23, all *p* < 0.0024; **Fig. 6D**). Interestingly, we found that the mean connectivity strength of selected links was positively predictive of inter-trial neural distinctiveness of the final run in the MTL (r = 0.39, *p* < 0.01; **Fig. 6E**). Taken together, these results indicate that retrieval-induced dynamic reconfiguration of memory-related brain networks is predictive of false memory and inter-trial distinctiveness in the MTL, with the most prominent effect in the MPFC.

**Fig. 6.**
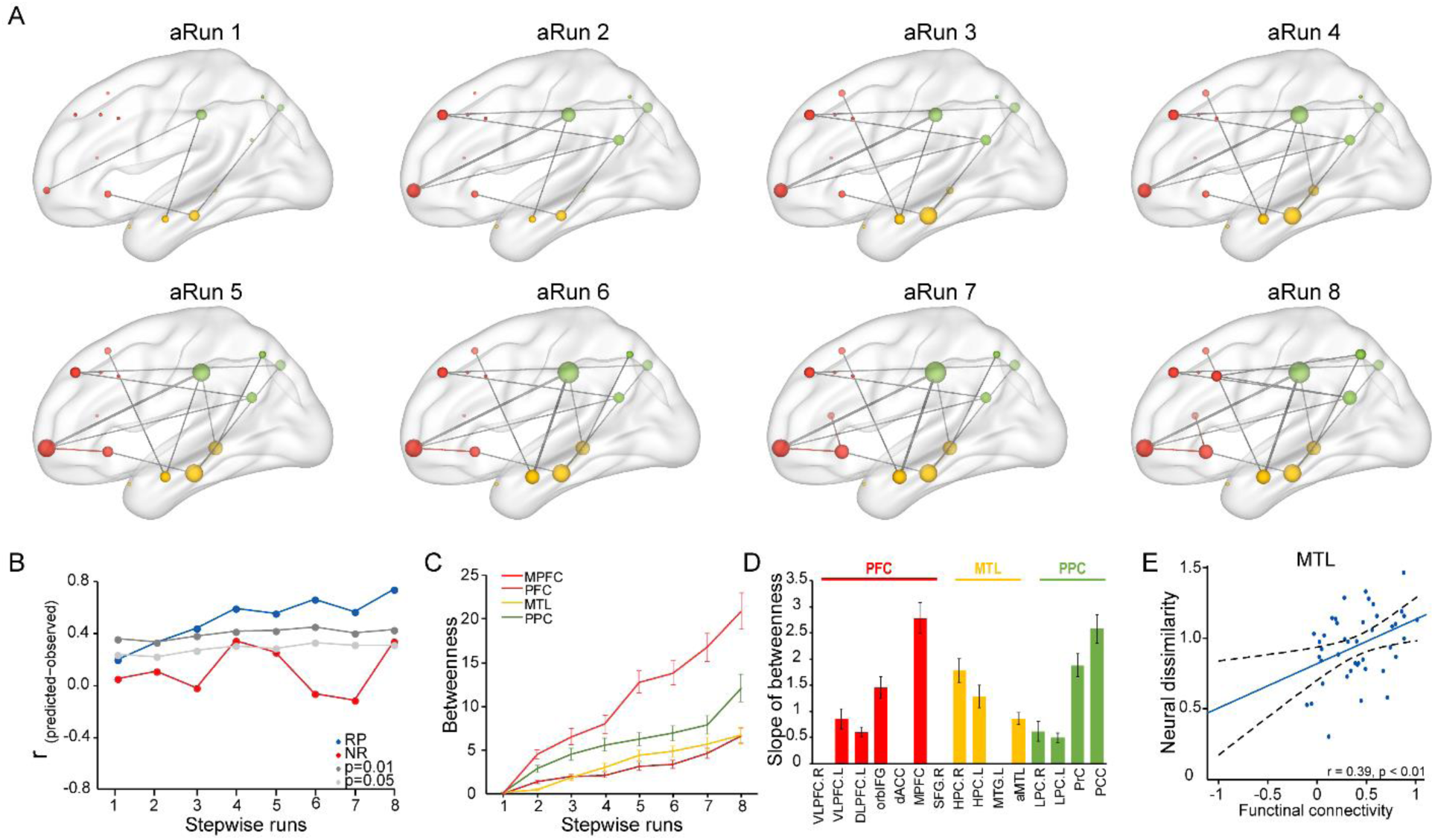
Brain network-based prediction of general false memory and network reconfiguration over retrieval practice. **(A)** Lateral views of selected links critical for predicting false memory outcomes project onto a glass brain to illustrate dynamic changes in memory-related neural network configurations over the course of 8 runs retrieval practice. Red, green and yellow nodes in the prefrontal cortex (PFC), medial temporal lobe (MTL) and posterior parietal cortex (PPC), respectively. **(B)** Line plots show prediction values from stepwise brain-behavior prediction analyses for false memory outcomes. Prediction values from the RP condition reach as high as 0.55 over the first five runs, and increase up to 0.74 across all 8 runs based on stepwise accumulative information as the progression of retrieval practice, which significantly outperforms the randomly-permutated level, and the NR condition. (**C**) Line plots show betweenness centrality in the medial prefrontal cortex (MPFC), MTL, PFC and PPC systems. (**D**) Bars depict the slopes of dynamic changes in betweenness centrality over 8 runs in 15 nodes for the RP condition. (**E**) Scatter plot shows a positive correlation between averaged functional connectivity of selected links and inter-trial neural distinctiveness of the MTL in the final run for the RP condition. Notes: Abbreviations for 15 nodes are listed in Supplemental Texts. Error bars represent standard error of mean.

## Discussion

By tracking dynamic changes in neural representations and network configurations, we investigated the neurocognitive mechanisms regarding how memories are reshaped by retrieval practice and subsequent consolidation. Retrieval practice boosted long-term retention after consolidation with a period of 24-hour but not a 30-minute interval, and increased false memory in general. These behavioral effects were associated with retrieval-induced dynamic changes in memory-related multi-voxel neural patterns as well as gradual reconfiguration of functional connections among PPC, PFC and MTL nodes. The VLPFC emerged as a hub for links that were critical in predicting long-term retention gains and refining neural pattern distinctiveness in the PPC, whereas the MPFC showed the most prominent effect for the selected links that predicted false memory outcomes and neural distinctiveness in the MTL. Our findings suggest that retrieval practice progressively refines memory-related neural representations and reconfigures functional routes to not only promote long-term retention gains but also to produce memory distortions.

At a behavioral level, retrieval practice boosted long-term retention after 24 hours but not 30 minutes. This is in line with previous studies reporting the benefits of retrieval practice on long-term but not immediate memory^1,3^. This suggests that the benefits of retrieval practice on memory are time-dependent, involving offline consolidation that transforms the retrieved memories into long-term store. We speculate that multiple retrieval attempts on the targeted memory traces may prioritize or “tagged” these memories as important or future-relevant, thus tune them into a super-ordinate position for long-term storage. This view can readily account for our observations on associative memory gains (for the important central information) while false memories for less important details. However, memories that did not go through retrieval practice are likely to set as “less important”, and thus would be forgotten over time. Sleep is known to be crucial for offline consolidation^32^. It is thus possible that processes during sleep may play a role in prioritizing the “tagged” information for subsequent consolidation^20^, thereby resulting in superior long-term retention gains.

It is worth noting that we observed a similar pattern of long-term retention gains for the negative and neutral scenes associated with faces. Similar to previous studies^33^, our observed benefit could be generalizable across both types of stimuli. In addition, our NR manipulation was seemingly analogous to memory suppression task^22,34^. However, we did not observe suppression-induced forgetting on a behavioral level. We would thus be cautious to regard this condition as voluntary suppression, given that our priori hypothesis in retrieval practice was with different experimental settings. One may conjecture the possibility of retrieval-induced forgetting^12^ that could favor long-term retention gains in the RP but not NR condition. Our data appears to speak against this assumption, because we did not observe retrieval-induced forgetting, but a prominent gain in long-term retention for RP condition in the delayed test. Future studies with memory suppression and retrieval-induced forgetting paradigms may directly address these possibilities.

Interestingly, retrieval practice also led to the recall of more illusory episodic details at both 24-hour and 30-min intervals for the RP condition as compared to the NR condition. This seemingly unwanted effect of retrieval practice is consistent with previous reports that retrieval increases the likelihood of false memory^7,35^. This is also in line with retrieval-induced learning models, suggesting that retrieving an episodic memory initiates new encoding, leaving the original memory trace vulnerable to being altered and thus supporting ever changing environmental needs^6,14^. Our observation of more false details recalled already in the immediate test suggests that retrieval practice can rapidly reshape original memories.

On the neural level, retrieval practice led to dynamic changes in multi-voxel activity patterns in memory-related brain systems, with the most prominent increase in inter-trial neural dissimilarity selectively seen in the PPC. The dissimilarity measure is recognized as a reliable metric of fine-tuned discriminative neural representations among individual traces in memory space^36^. This finding likely signifies that memory representations for the retrieved events become increasingly differentiated from each other in the PPC over retrieval practice. In other words, representations of individual memories become more discrete in memory space to better support access to each individual memory^11^. Critically, inter-trial neural distinctiveness in the final run in the RP but not NR condition was positively predictive of long-term retention gains after consolidation. This association did not emerge in the first run, suggesting the evolvement of fine-tuned neural representations is a progressive process over retrieval practice. In fact, a recent study revealed that memory engrams can be formed in the PPC within hours^18^. It is possible to assume that PPC may play a role in tuning memory representations into a differentiated status over retrieval practice, and offline consolidation across the 24-hour delay, likely during sleep^37^, may further stabilize these representations for long-term store.

Interestingly, we found that inter-trial neural distinctiveness for the RP but not NR trials was predictive of false memory in the MTL. This is in line with previous findings that the MTL especially the hippocampus contributes to false memory^38^. The MTL is crucial for episodic memory, including episodic context binding and recollection^39^. Episodic contexts may serve as important cues to support the benefits of retrieval practice^9^, but mismatch of episodic contexts can lead to false memory in humans^40^. Thus, the MTL could be involved in online reinstatement of memories in neocortical ensembles, through which memories become labile and vulnerable to alteration during retrieval practice. By this view, inter-trial neural distinctiveness in the MTL is responsible for the generation of false memory, most likely due to retrieval-mediated learning of new information and/or mismatching episodic context during multiple retrieval attempts.

Beyond retrieval-induced changes in local neural representations, we found that functional connectivity among large-scale brain networks became strengthened over 8 runs, with higher global network efficiency in the RP than NR condition. Inter-regional connectivity is thought to reflect functional communication routes of information flow between distant regions^41^. The strengthening of connectivity with higher efficiency among memory-related brain networks may reflect a gradual build-up of more effective routes during retrieval practice, through which the target information could be reweighted as important for future use. Moreover, our network-based prediction results show that retrieval-induced dynamic reconfiguration of brain networks over 8 runs is critical to predict long-term retention gains, with a rapid growth in betweenness centrality during the initial five runs. And the most prominent effect of this emerged in the right VLPFC. As betweenness reflects the transfer of information flow through a network^42^, the VLPFC may thus act as a hub to drive dynamic network reconfiguration over retrieval practice. The MTL exhibited a generally low yet similar dynamic trajectory to the PFC and PPC, indicating a constant engagement of these systems in retrieval practice. Given long-term retention gains observed after offline consolidation, we speculate that the above network reconfiguration may be preparatory for setting up relevant connections to be prioritized for subsequent consolidation into long-term store.

We further found that the right VLPFC connections with the PPC during retrieval practice provided the most information to predict long-term retention gains after consolidation. The VLPFC is recognized as a key locus that exerts top-down modulation of episodic retrieval, involving strategy selection^24^, and suppression of irrelevant information^43^. The PPC has been considered as an “output buffer” for active representations of retrieved memory content, through converging multimodal information from other cortical inputs^29,44^. We thus speculate that the gradually strengthened connectivity of the VLPFC with PPC might underlie efficient functional coordination to facilitate reinstatement of the target memory representations anchored in the PPC over retrieval practice. With repeatedly active retrieval, the routes between these nodes are gradually established, and the target information can thus be reweighted as high priority for consolidation into long-term store. Indeed, we observed that the VLPFC-PCC functional connectivity was positively predictive of inter-trial neural distinctiveness in the PPC, and more discriminative neural patterns in the PPC were further predictive of better long-term retention gains after consolidation. Recent rodent models posit that the formation of long-term memory involves early ‘tagging’ and reweighting of cortical networks that subsequently support the memory^45,46^. It is thus possible that the VLPFC could drive an evaluation signal to reweight the repeatedly reactivated memory events as important over retrieval practice, through which representations of the target information in the PPC might be prioritized into a super-ordinate position for subsequent consolidation. This prefrontal modulation differs from the top-down control of the dorsolateral PFC over neural activity in the hippocampus implicated in voluntary suppression of memories^22,34^. Critically, the reweighted memory traces during retrieval practice appeared to mature through the process of offline consolidation, as we did not observe memory gains in the immediate recall, which might due to the possibility that they are still labile and not matured into the stabilized engrams^18^, and not completely segregated from others during immediate recall. Indeed, recent data suggests that sleep is critical to preserve newly formed memory engrams^37^. Further investigations are needed to determine the mechanisms of how memory traces are prioritized through VLPFC and PPC coordination over retrieval practice to foster later consolidation.

In conjunction with neural network reorganization to predict long-term retention gains, we found that reconfiguration of memory-related neural networks over retrieval practice could predict false memory outcomes. Unlike the VLPFC and PPC, which appeared critical for predicting long-term retention gains, we found the most prominent increases in betweenness in the MPFC and PCC. This suggests that functional coordination between the MPFC and PCC over retrieval practice plays a critical role in producing false memory. Such interpretation is in line with previous findings on the constructive nature of episodic memory systems. Namely, the MPFC and PCC may act as an interface for scaffolding active reconstruction of episodic memories for future use, and thereby leads to retrieval-induced new learning to produce false memory in subsequent recall^47^. In addition, these nodes are known to support recall of autographical memory^48^, which might interfere target memories by integrating one’s autographical memories.

Finally, we found that the mean connectivity strength across selected links was positively predictive of inter-trial neural distinctiveness of the MTL in the final run, and the latter measure further correlated with more false memory. The MTL has been implicated in constructive processes for online reinstatement of episodic memories during retrieval practice, through which memory representations become labile at each retrieval and thus new information of the current experience is likely to be integrated into existing memories leading memory representations labile at each retrieval^49^. This expands the central idea of research on the benefits of retrieval practice on memory retention, by highlighting that retrieval practice reshapes memories through dynamic reorganization of neural representations and network connectivity to produce false memories. Altogether, our findings provide new evidence to suggest distinct retrieval-induced dynamic neural reorganization mechanisms through which retrieval practice can lead to seemly contradicting memory outcomes including on the one hand long-term retention gains after consolidation and on the other hand, false memory in general.

### In conclusion

our study demonstrates that rapid neural reorganization during retrieval practice predicts long-term retention gains after consolidation and false memory outcomes. Our findings suggest a dynamic neural reorganization mechanism, through which memories are arranged into more discrete yet malleable representations via rapid reorganization of brain functional routes for subsequent consolidation into long-term memory. Our findings may inform the development of effective interventions and strategies to improve memory in both healthy and clinical populations.

## Materials and Methods

### Participants

Fifty-seven young, healthy college students (32 females, average age ± s.d.: 22.37 ± 2.22 years old, range from 19 to 29) participated in this study, and had intact associative memory data. All participants were right-handed with normal or corrected-to-normal vision, and reported no history of neurological or psychiatric disease. The Institutional Review Board approved the study protocol for Human Subjects at Beijing Normal University, and written informed consent was obtained from all participants before the experiment. However, due to unexpected data unavailability the sample size of false memory reduced to fifty. Data from seven participants were excluded from further analyses due to excessive head motion during fMRI scanning with root mean squared motion parameters over a voxel’s width. To sum up, this resulted in 56 participants for associative memory and 50 participants for false memory as the behavioral sample size, with 50 and 43 as their corresponding imaging data sample size, respectively.

### Materials

Forty-eight pairs of face-scene associations were used in this study. Faces with neutral expressions were selected from a standardized Chinese face database^22^. Briefly, faces were carefully selected using the following criteria: direct gaze contact, no obvious emotional facial expression, no headdress, no glasses and no beard. Forty-eight complex scenes were selected from the International Affective Picture System (IAPS). Neutral scenes were selected with a modest level of emotional valence (mean ± s.d.: 5.32 ± 0.47) and low level of arousal (mean ± s.d.: 2.52 ± 0.36) as measured on a 9 - point scale (1 = ‘extremely negative’ or ‘not arousing at all’, 9 = ‘extremely happy’ or ‘extremely arousing). Negative scenes were selected with negative emotional valence (mean ± s.d.: 2.37 ± 0.69) and high level of arousal (mean ± s.d.: 7.89 ± 0.55). Faces and scenes had been used in an independent study with a minimal relatedness in content to each other, and matched on luminance^22^. Faces and scenes were randomly paired to create 48 face-scene associations across participants.

### Experimental design and procedure

The experiment consisted of three phases: memory acquisition, retrieval practice, and two subsequent memory tests after 30-min and 24-h intervals respectively. In the memory acquisition phase outside the scanner, participants were trained to memorize 48 face-scene associations with multiple study-test cycles. For each cycle, each face-scene association was presented for 6 seconds in the center of the screen, and participants were instructed to remember face-scene associations for subsequent memory tests. Thereafter, they performed an associative memory recognition test in which a given face in the upper center of the screen and two scenes on the left and right of the lower screen were displayed. One scene was the correct association while the other one randomly selected from other associations. Participants were asked to select either the left or the right scene as the picture associated with that face. There was no feedback given either by the experimenters or the task program. After a cycle of associative recognition tests was finished, the program immediately scored the participant’s performance. The training session would continue until the participant reached at least 90% accuracy.

During the memory practice phase inside the scanner, 32 face-scene associations were randomly selected from the acquisition phase and the remaining 16 associations served as the baseline condition and were not presented in this phase. For each trial, a face surrounded by either a blue or red rectangle frame was presented for 4 seconds. Participants were instructed to engage in either active retrieval practice (i.e., RP, 16 trials, cued with a blue rectangle) or passively view the face without trying to retrieve the associated scene (NR, 16 trials, cued with a red rectangle) of the scene associated with the face cue. We chose the NR condition as a dedicated control condition with matched visual stimulation. Thereafter, participants were asked to rate the vividness of recalled memories on a four-point scale (1 = ‘Not at all; 4 = ‘Extremely’)^1^. The original scene was not shown again, in accordance with many studies on retrieval practice without feedback^1,3,27^. Trials were jittered with an inter-trial interval varying from 2 to 6 seconds (average = 4 seconds with 1 second step). The entire memory practice phase consisted of 8 runs with two-minute breaks between runs, and lasted 36 minutes in total, with 4.5 minutes for each run.

In the memory test phase outside the scanner, memory performance for face-scene associations was assessed by two independent cued-recall tests: one was performed after 30 minutes (i.e., immediate recall test), and the other one was performed after 24 hours (i.e., delayed recall test). All 32 faces (i.e., 16 from the RP, and 16 from the NR condition) from the retrieval practice phase were randomly split into two halves as cues for the immediate and delayed recall tests respectively. Participants were asked to orally recall and describe the scenes associated of each face cue. It is worth noting that 16 face-scene associations from the memory acquisition phase did not appear during the retrieval practice phase, and were also split into two halves corresponding to immediate and delayed recall tests. The presentation of face cues was randomized at test across participants. For each face cue, participants had a maximum of 30 seconds to verbally describe the associated scene with audio recording. Two raters who were blind to the purposes of the study scored each participant’s oral recall independently. When any inconsistences were encountered, a final score of each item was made by further discussion with a final consensus between the raters. All participants reported adequate sleep during the night after the retrieval practice, with approximately 8 hours of sleep (average 7.68 ± 1.05 hours).

### Memory performance and behavioral analyses

To assess episodic memory for face-scene associations, the raters scored participants’ memory performance according to their oral recall of the information that was enough to identify the associated scenes, as well as false information recalled (e.g., a participant recalled “a blue book” as “a black book”). Each participant’s memory accuracy (correct proportion) was quantified as the amount of face-scene associations later remembered as a proportion from all associations within each condition (RP, NR, baseline) and testing (immediate/30-minute, delayed/24-hour). Each participant’s false memory was quantified as the sum of false information recalled for the complex scenes associated with each face that were later remembered for the associative memory. We further computed the memory retention score for the immediate and delayed tests by subtracting the memory accuracy from the corresponding baseline condition to provide a measure of retrieval practice efficiency. This retention measure was then submitted to a 2-by-2 repeated measure of ANOVA with Condition (RP vs. NR) and Interval (Immediate vs. Delayed) as within-subject factors. Each participant’s long-term retention gain after consolidation was computed by subtracting retention scores in the immediate recall test from that of the delayed recall test. Pearson’s correlations and prediction analyses were conducted to assess the relationship of retrieval-induced changes in neural representations and network reconfiguration with long-term retention gains and false memory outcomes.

### Imaging acquisition

Whole-brain images were acquired on a Siemens Trio 3.0 Tesla MR scanner. Functional images were acquired using an echo-planar imaging sequence (37 slices; TR, 2000ms; TE, 30ms; flip angle, 90°; voxel size, 3.5 × 3.5 × 3.5mm; FOV, 224 × 224mm). High-resolution T1-weighted anatomical images were acquired by using a magnetization-prepared rapid acquisition gradient echo (MP-RAGE) sequence (144 slices; TR, 2530ms; TE, 3.39ms; voxel size, 1.3 × 1.0 × 1.3mm; flip angle, 7°; FOV, 256 × 256mm).

### Imaging preprocessing

Brain images were preprocessed using Statistical Parametric Mapping toolbox (SPM8; http://www.fil.ion.ucl.ac.uk/spm). The first 4 volumes of functional images were discarded to allow for signal equilibrium. Remaining images were realigned to the mean image of each run and corrected for slice acquisition timing. Subsequently, functional images were co-registered to each participant’s gray matter image segmented from corresponding T1-weighted image and spatially normalized into the stereotactic template of the Montreal Neurological Institute (MNI). Finally, images were smoothed using a 6-mm FWHM Gaussian kernel.

### Univariate GLM analysis

To assess task-related brain responses during memory practice, separate regressors of interest were modeled for trials in the RP and NR conditions, and convolved with the canonical hemodynamic response function (HRF) in SPM8. In addition, each participant’s motion parameters from the realignment procedure were included to regress out potential effects of head movement on brain response. We included high-pass filtering using a cutoff of 1/128 hz to remove high frequency noise and corrections for serial correlations using a first-order autoregressive model (AR(1)) in the GLM framework.

Contrast parameter estimate images for task-related brain responses in RP (or NR) condition versus lower-level fixation, generated at the individual-subject level, were submitted to subsequent analyses for multi-voxel representation stability and fidelity as well as graph theory-based brain network analyses over the course of retrieval practice.

To further assess trial-wise brain responses for each RP or NR trial during the memory practice phase, each cue run was modeled as a separate regressor and convolved with the HRF implemented in SPM8. This resulted in a total of 32 regressors for each run, with 16 RP trials and 16 NR trials. The other parameter settings were the same as above univariate GLM for task-related estimation of brain responses. Contrast parameter estimate images for each retrieved trial versus fixation, initially generated at the individual-subject level, were submitted to subsequent analyses for inter-trial multi-voxel pattern distinctiveness and trial-specific neural signatures over 8 runs in RP and NR conditions separately.

### ROI Selection

To define a memory-related brain mask, we used the Neurosynth platform for large-scale, automated synthesis of fMRI data (http://neurosynth.org) with ‘memory retrieval’ as a search term and generated a reverse inference mask at the whole-brain level. We then refined the mask using a criterion of a height threshold of *p* < 0.001 (or z-score > 3.0) and a spatial extent cluster size of more than 30 voxels. Then, the whole-brain mask was segmented into 15 different ROIs or clusters based on spatially contiguous voxels. These ROIs included the left ventral lateral prefrontal cortex (l_VLPFC), right ventral lateral prefrontal cortex (r_VLPFC), left dorsal lateral prefrontal cortex (l_DLPFC), orbito-inferior frontal gyrus (orbIFG), dorsal anterior cingulate cortex (dACC), medial prefrontal cortex (MPFC), right superior frontal cortex (r_SFG), right hippocampus (r_HPC), left hippocampus (l_HPC), left medial temporal gyrus (l_MTG), anterior medial temporal lobe (aMTL), right lateral parietal cortex (r_LPC), left lateral parietal cortex (l_LPC), precuneus (PrC) and posterior cingulate cortex (PCC).

### Neural pattern similarity and fidelity

Three multi-voxel neural pattern metrics were computed to characterize changes in retrieval-induced multi-voxel pattern similarity over the course of retrieval practice. The first one referred to neural pattern similarity over 8 runs during retrieval practice. The z-score map of retrieval-induced neural activation pattern was first obtained from the Neurosynth platform as a canonical reference by using the overall brain mask defined above. Multi-voxel activity patterns for RP and NR conditions were then separately extracted from the same mask in each run. And then we computed Pearson’s correlation coefficients for the respective multi-voxel patterns of RP and NR with the canonical reference map for each run. Thereafter, correlation coefficients were Fisher’s z-transformed and submitted to compute neural representation stability S_similarity across 8 runs for RP and NR separately.

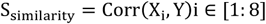

where X_i_ is condition-related activity patterns of run i for each condition, and Y is multi-voxel activity pattern from the Neurosynth.

The other two metrics, reflecting condition-related neural fidelity and trial-specific neural pattern similarity over retrieval practice, were described in the Supplemental Methods.

### Inter-trial neural pattern distinctiveness

A neural pattern distinctiveness metric was computed to characterize changes in inter-trial multi-voxel activity pattern dissimilarity among RP (or NR) trials over the course of 8-run retrieval practice. The multi-voxel activity pattern for each trial **i**n the RP (or NR) condition was extracted from the mask in each run. We then computed their corresponding Pearson’s correlation coefficients with activity patterns of other remaining trials from the same condition of the same run. These coefficients were transformed into Fisher’s z-scores and averaged, and the average was then subtracted from 1 to yield a distinctiveness metric for each run. We then fitted the linear equation function [f(x) = a × x + b, where variable x is the run number] to the dynamic neural pattern distinctiveness.

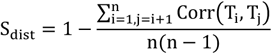

where n is the total number of trials in each condition, T_i_ is trial-related activity pattern associated with trial i and T_j_ is trial-related activity pattern for trial j from the same condition of the run. Due to the fact that the last run (the 8^th^ run in this study) represents the final brain state after memory practice, we used the 8^th^ run for further correlational analyses with memory performance as implemented by previous studies^24^.

### Network construction

Network nodes consisted of 15 ROIs as defined above. For each participant, a 15 × 15 connectivity matrix was created for each task condition (i.e., RP or NR) in each run by using a generalized form of context-dependent psychophysiological interaction (gPPI) analysis. The gPPI approach was widely used to assess task-dependent functional connectivity of a specific seed or ROI with the rest of the brain, after removing potential confounds of overall task activation and common driving inputs. Specifically, mean time series from each seed ROI were extracted and then deconvolved so as to uncover neuronal activity (i.e., physiological variable) and multiplied with the task design vector contrasting the RP condition versus the fixation condition (i.e., a binary psychological variable) to form a psychophysiological interaction (PPI) vector. This interaction vector was convolved with a canonical HRF in SPM8 to form the PPI regressor of interest. The psychological variable representing task design (RP versus fixation) as well as mean-corrected time series of each seed ROI were also included in the GLM to remove overall task-related activation and the effects of common driving inputs on brain connectivity. To ensure normality, connectivity values of each task condition were Fisher’s z-transformed. Note that only the voxels within 15 ROIs were included in this analysis to save computational resources. Separate gPPI analyses were conducted for each seed ROI to assess its task-dependent functional connectivity with the remaining ROIs.

### Network-based brain-behavior prediction analysis

Network matrices derived from the gPPI analyses were submitted to subsequent brain-behavior prediction analyses based on machine learning algorithms using the LIBSVM toolbox (http://www.csie.ntu.edu.tw/~cjlin/libsvm/). In the machine learning-based prediction model, a leave-one-out approach was used to train separate models to predict individual’s long-term retention gains after consolidation as well as false memory outcomes. To predict an individual’s long-term retention gain, for instance, we trained a Support Vector Regression (SVR) model with the data of remaining individuals. This approach was iterated to compute predicted long-term retention scores for all individuals. We then calculated Pearson’s correlation coefficients between predicted and observed scores of the prediction accuracy to quantify the strength of the brain-behavior relationship. The statistical significance of prediction accuracy was assessed by a permutation test procedure. For each permutation, we randomly shuffled behavioral scores, and then computed the brain-behavior prediction accuracy. This procedure was iterated for 1,000 times for each run. Thereafter, 1000 permutated prediction values were sorted in a descending order, and the significance p values of 0.01 and 0.05 were computed by dividing the position number (that the prediction accuracy located) by 1000 for each run.

### Two-step prediction procedures were used to compute brain-behavior prediction values

First, we pulled all links across 8 runs as input features to train a SVR model in order to predict an individual’s long-term retention gain (or false memory score). In this model, we obtained the predictive weight for each link, which we sorted in descending order. Subsequently, only links whose weights were above certain criteria were selected, and used as input features in the model for the follow-up stepwise prediction procedure. Following the convention in the field, we implemented a set of different thresholds and obtained the top 1% of links to achieve the best prediction accuracy (**Fig. S7A**). Second, we then projected those top 1% links back to each run to track their evolution trajectories over the progression of 8 runs in a cumulative way, and then trained a stepwise prediction model to predict individual’s long-term retention gains (or false memory scores) (Fig. 4A). Furthermore, these selected links were submitted for graph theoretical analyses and network visualization by superimposing onto a glass brain template.

### Global efficiency

Global efficiency represents the average inverse shortest path length in a network, and is inversely related to the characteristic path length. Global efficiency is defined as

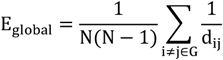

where G is the network graph, i and j are ROIs, d_ij_ is a measure of distance between i and j (obtained from Euclidean distance).

### Betweenness

As one of the most frequently used metrics in network analysis, betweenness centrality represents the degree to which information pass through a node, the more of which, the higher influence of this node has within a network^42^. It was computed according to the networks with selected features from prediction analysis. Specifically, links that were predictive of individual’s long-term retention gains after consolidation were maintained and the remaining ones were set as 0. Thereafter, the resultant networks were entered into GRETNA for network analysis (https://www.nitrc.org/projects/gretna/). The betweenness centrality of node i was defined as follows:

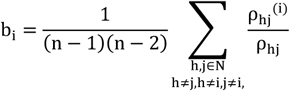

Where N is the set of all nodes in the network, and n is the number of nodes. ρ_hj_is the number of shortest paths between h and j, and ρ_hj_^(i)^ is the number of shortest paths between h andj that pass through i.

### Statistical analysis

Statistical testing for behavioral and imaging data was performed using R(3.4.1) and MATLAB(R2016a) respectively. Values are presented as mean ± s.e.m., unless indicated otherwise. Repeated ANOVAs and Student’s t-tests were used to assess differences between conditions of interest for equal variances, when normality was assumed. Pearson’s correlation and linear regression were used to assess the relationship between variables and linear fitting when appropriate. The two-tailed *p* values are reported for statistical testing. The permutation test was implemented for the prediction analyses.

### Data availability

The data and codes that support the findings of this study are available from the corresponding author on request.

## Acknowledgements

We thank Yunzhe Liu and Wanjun Lin for their assistance in conducting the experiment and data collection. We also thank Frederik D. Weber for his valuable comments for the manuscript.

## Author contributions

S.Q. conceived the experiment. B.X. performed data collection. L.Z. J.W., C.B. and H.L. performed data analysis. L.Z., J.W., P.J.B. and S.Q. wrote the manuscript. All authors contributed to data discussion and interpretation.

## References

1 Karpicke, J. D. & Roediger, H. L., 3rd. The critical importance of retrieval for learning. Science 319, 966–968, doi: 10.1126/science.1152408 (2008).

2 Racsmány, M., Conway, M. & Demeter, G. Consolidation of Episodic Memories During Sleep. Psychol. Sci. 21, 80–85 (2010).

3 Roediger, H. L., 3rd & Butler, A. C. The critical role of retrieval practice in long-term retention. Trends in cognitive sciences 15, 20–27, doi: 10.1016/j.tics.2010.09.003 (2011).

4 Pastötter, B. & Bäuml, K.-H. T. Testing enhances subsequent learning in older adults. Psychology and aging 34, 242 (2019).

5 Gershman, S. J., Monfils, M.-H., Norman, K. A. & Niv, Y. The computational nature of memory modification. Elife 6, e23763 (2017).

6 Nadel, L. & Land, C. Commentary—reconsolidation: memory traces revisited. Nat. Rev. Neurosci. 1, 209 (2000).

7 Roediger III, H. L., Jacoby, J. D. & McDermott, K. B. Misinformation effects in recall: Creating false memories through repeated retrieval. Journal of Memory and Language 35, 300–318 (1996).

8 Van den Broek, G. et al. Neurocognitive mechanisms of the “testing effect”: A review. Trends Neurosci. Educ. 5, 52–66, doi: 10.1016/j.tine.2016.05.001 (2016).

9 Karpicke, J. D., Lehman, M. & Aue, W. R. in Psychology of learning and motivation Vol. 61 237–284 (Elsevier, 2014).

10 Antony, J. W., Ferreira, C. S., Norman, K. A. & Wimber, M. Retrieval as a Fast Route to Memory Consolidation. Trends in cognitive sciences 21, 573–576, doi: 10.1016/j.tics.2017.05.001 (2017).

11 Wimber, M., Alink, A., Charest, I., Kriegeskorte, N. & Anderson, M. C. Retrieval induces adaptive forgetting of competing memories via cortical pattern suppression. Nat. Neurosci. 18, 582–589, doi: 10.1038/nn.3973 (2015).

12 Hulbert, J. C. & Norman, K. A. Neural Differentiation Tracks Improved Recall of Competing Memories Following Interleaved Study and Retrieval Practice. Cereb. Cortex 25, 3994–4008, doi: 10.1093/cercor/bhu284 (2015).

13 McDaniel, M. A. & Masson, M. E. Altering memory representations through retrieval. J. Exp. Psychol. Learn. Mem. Cogn. 11, 371 (1985).

14 Yassa, M. A. & Reagh, Z. M. Competitive trace theory: a role for the hippocampus in contextual interference during retrieval. Front. Behav. Neurosci. 7, 107 (2013).

15 Epp, J. R., Mera, R. S., Köhler, S., Josselyn, S. A. & Frankland, P. W. Neurogenesis-mediated forgetting minimizes proactive interference. Nature communications 7, 10838 (2016).

16 McClelland, J. L., McNaughton, B. L. & O’Reilly, R. C. Why there are complementary learning systems in the hippocampus and neocortex: insights from the successes and failures of connectionist models of learning and memory. Psychol. Rev. 102, 419 (1995).

17 Tonegawa, S., Morrissey, M. D. & Kitamura, T. The role of engram cells in the systems consolidation of memory. Nat. Rev. Neurosci., 1 (2018).

18 Brodt, S. et al. Fast track to the neocortex: A memory engram in the posterior parietal cortex. Science 362, 1045 (2018).

19 Yang, G. et al. Sleep promotes branch-specific formation of dendritic spines after learning. Science 344, 1173–1178, doi: 10.1126/science.1249098 (2014).

20 Stickgold, R. & Walker, M. P. Sleep-dependent memory triage: evolving generalization through selective processing. Nat. Neurosci. 16, 139 (2013).

21 Redondo, R. L. & Morris, R. G. Making memories last: the synaptic tagging and capture hypothesis. Nat. Rev. Neurosci. 12, 17 (2011).

22 Liu, Y. et al. Memory consolidation reconfigures neural pathways involved in the suppression of emotional memories. Nat. Commun. 7, 13375, doi: 10.1038/ncomms13375 (2016).

23 Born, J. & Wilhelm, I. System consolidation of memory during sleep. Psychol Res 76, 192–203, doi: 10.1007/s00426-011-0335-6 (2012).

24 Kuhl, B. A., Dudukovic, N. M., Kahn, I. & Wagner, A. D. Decreased demands on cognitive control reveal the neural processing benefits of forgetting. Nat. Neurosci. 10, 908 (2007).

25 Hutchinson, J. B., Uncapher, M. R. & Wagner, A. D. Posterior parietal cortex and episodic retrieval: convergent and divergent effects of attention and memory. Learning & Memory 16, 343–356 (2009).

26 Kriegeskorte, N. & Kievit, R. A. Representational geometry: integrating cognition, computation, and the brain. Trends in cognitive sciences 17, 401–412, doi: 10.1016/j.tics.2013.06.007 (2013).

27 Kuhl, B. A., Shah, A. T., DuBrow, S. & Wagner, A. D. Resistance to forgetting associated with hippocampus-mediated reactivation during new learning. Nat. Neurosci. 13, 501–506, doi: 10.1038/nn.2498 (2010).

28 Ranganath, C. et al. Dissociable correlates of recollection and familiarity within the medial temporal lobes. Neuropsychologia 42, 2–13 (2004).

29 Wagner, A. D., Shannon, B. J., Kahn, I. & Buckner, R. L. Parietal lobe contributions to episodic memory retrieval. Trends in cognitive sciences 9, 445–453, doi: 10.1016/j.tics.2005.07.001 (2005).

30 Bassett, D. S., Yang, M., Wymbs, N. F. & Grafton, S. T. Learning-induced autonomy of sensorimotor systems. Nat. Neurosci. 18, 744–751, doi: 10.1038/nn.3993 (2015).

31 Dresler, M. et al. Mnemonic Training Reshapes Brain Networks to Support Superior Memory. Neuron 93, 1227–1235 e1226, doi: 10.1016/j.neuron.2017.02.003 (2017).

32 Diekelmann, S. & Born, J. The memory function of sleep. Nat. Rev. Neurosci. 11, 114 (2010).

33 Jia, X., Gao, C., Cui, L. & Guo, C. Does emotion arousal influence the benefit received from testing: insights from neural correlates of retrieval mode effect. Neuroreport 29, 1449–1455, doi: 10.1097/wnr.0000000000001130 (2018).

34 Anderson, M. C. et al. Neural systems underlying the suppression of unwanted memories. Science 303, 232–235, doi: 10.1126/science.1089504 (2004).

35 Mcdermott, K. B. Paradoxical effects of testing: Repeated retrieval attempts enhance the likelihood of later accurate and false recall. Mem. Cognit. 34, 261–267 (2006).

36 Wirebring, L. K. et al. Lesser neural pattern similarity across repeated tests is associated with better long-term memory retention. J. Neurosci. 35, 9595–9602 (2015).

37 Himmer, L., Schönauer, M., Heib, D. P. J., Schabus, M. & Gais, S. Rehearsal initiates systems memory consolidation, sleep makes it last. Science advances 5, eaav1695 (2019).

38 Schacter, D. L. & Slotnick, S. D. The cognitive neuroscience of memory distortion. Neuron 44, 149–160 (2004).

39 Jeong, W., Chung, C. K. & Kim, J. S. Episodic memory in aspects of large-scale brain networks. Front Hum Neurosci 9, 454, doi: 10.3389/fnhum.2015.00454 (2015).

40 Loftus, E. F. Planting misinformation in the human mind: a 30-year investigation of the malleability of memory. Learn Mem 12, 361–366, doi: 10.1101/lm.94705 (2005).

41 Tanimizu, T. et al. Functional Connectivity of Multiple Brain Regions Required for the Consolidation of Social Recognition Memory. J. Neurosci. 37, 4103–4116, doi: 10.1523/JNEUROSCI.3451-16.2017 (2017).

42 Freeman, L. C. A set of measures of centrality based on betweenness. Sociometry, 35–41 (1977).

43 Guo, Y., Schmitz, T. W., Mur, M., Ferreira, C. S. & Anderson, M. C. A supramodal role of the basal ganglia in memory and motor inhibition: Meta-analytic evidence. Neuropsychologia 108, 117–134, doi: https://doi.org/10.1016/j.neuropsychologia.2017.11.033 (2018).

44 Sestieri, C., Shulman, G. L. & Corbetta, M. The contribution of the human posterior parietal cortex to episodic memory. Nature reviews. Neuroscience 18, 183–192, doi: 10.1038/nrn.2017.6 (2017).

45 Tonegawa, S., Morrissey, M. D. & Kitamura, T. The role of engram cells in the systems consolidation of memory. Nature reviews. Neuroscience 19, 485–498, doi: 10.1038/s41583-018-0031-2 (2018).

46 Lesburguères, E. et al. Early Tagging of Cortical Networks Is Required for the Formation of Enduring Associative Memory. Science 331, 924, doi: 10.1126/science.1196164 (2011).

47 Warren, D. E., Jones, S. H., Duff, M. C. & Tranel, D. False Recall Is Reduced by Damage to the Ventromedial Prefrontal Cortex: Implications for Understanding the Neural Correlates of Schematic Memory. The Journal of Neuroscience 34, 7677, doi: 10.1523/JNEUROSCI.0119-14.2014 (2014).

48 Maddock, R. J., Garrett, A. S. & Buonocore, M. H. Remembering familiar people: the posterior cingulate cortex and autobiographical memory retrieval. Neuroscience 104, 667–676, doi: 10.1016/s0306-4522(01)00108-7 (2001).

49 Barry, D. N. & Maguire, E. A. Remote Memory and the Hippocampus: A Constructive Critique. Trends in cognitive sciences 23, 128–142, doi: 10.1016/j.tics.2018.11.005 (2019).

