## Supplementary Materials for "Rapid Neural Reorganization during Retrieval Practice Predicts Subsequent Long-term Retention and False Memory"

**Supplementary Methods**

***Neural pattern fidelity.*** The other metric, reflecting neural representation fidelity over the course of retrieval practice, was used to quantify the similarity of multi-voxel neural activity patterns between the initial run and each of the remaining runs (2^nd^ to 8^th^). In other words, this measure reflects the degree to how multi-voxel activity pattern in each of other runs is similar to its initial state of the first run. It was calculated according to the formula:

$$S_{\mathrm{fidelity}}=\mathrm{Corr}\left( X_{1},Y_{i} \right)i\in[2:8]$$

where X1 and Yi are condition-specific neural activity patterns in its initial state and run i respectively for each condition.

***Trial-specific neural pattern similarity.*** Trial-specific neural pattern similarity was computed to characterize how trial-specific multi-voxel activity pattern changes over 8 runs in the RP and NR conditions. Multi-voxel activity pattern for each trial was first extracted separately from the RP and NR conditions in each run. We then computed Pearson's correlation coefficients between each trial's multi-voxel activity pattern in the first run and the same trial's multi-voxel activity pattern in each of the remaining runs. These coefficients were transformed into Fisher's z-scores and averaged across all trials within same condition in each run. After that, we fitted a linear regression model for this data to estimate dynamic changes in trial-specific neural pattern similarity as a function of 8 runs in the RP and NR conditions separately.

***Modular segregation.*** The participation coefficient, as a measure of network segregation was computed to quantify changes in functional segregation of memory-related brain networks over retrieval practice^1^. In general, participation coefficient measures the strength of a node’s connection in its community, which reflects the tightness of connections between different modules. A module that has a high participation coefficient tends to display stronger functional communication within its community but less communication with other modules. The participation coefficient *Pi* of node *i* is defined as:

$$P_{i}=1-\sum_{m\in M} \left( \frac{k_{i}(m)}{k_{i}} \right)^{2}$$

where *m* is a module in a set of modules *M*, and *k_i_(m)* is the weight of connections between node *i* and all nodes in module *m*. To quantify the segregation of specific modules, we average *Pi* across all brain regions assigned to the same module. To quantify global network segregation, we average *Pi* across all nodes in the network.

**Supplementary Results**

**Vividness ratings during retrieval practice in relation to long-term retention gains and neural measures**

To examine whether vividness ratings during memory practice predict long-term memory gains, we conducted additional analyses for vividness ratings over 8 runs for RP and NR conditions and their relations to subsequent memory outcomes. A 2-by-8 repeated ANOVA with Run (1 to 8 runs) and Condition (RP vs. NR) as within-subject factors revealed a main effect of Condition (F_(1,56)_ = 258.78, p < 0.001), with significantly higher vividness ratings for the RP than NR condition in each run (follow-up paired t tests: all t_(56)_ > 14.34, all p<0.001) as well as on the average level (t_(56)_ = 24.93, p < 0.001) (**Fig. S1A&*C***). We further estimated the slope of each participant’s vividness ratings over 8 runs for the RP and NR conditions separately. We found a flat pattern of vividness ratings over 8 runs in the RP condition (beta = 0.011, t_(56)_ = 1.53, p = 0.13), but a gradual decrease in vividness ratings in the NR condition (beta = -0.022, t_(56)_ = -2.89, p = 0.006) (**Fig. S1*B***).

We observed a significant correlation of average vividness rating with false memory scores in RP (r = 0.38, p = 0.007), but not in NR condition (r = 0.20, p = 0.16). However, correlation analyses revealed no reliable correlations of average vividness ratings with long-term retention gains after consolidation in both RP (r = -0.09, p = 0.49) and NR conditions (r = -0.20, p = 0.13). Likewise, we did not observe any reliable correlations between the slope of vividness ratings over 8 runs with long-term retention gains after consolidation in both RP (r = -0.15, p = 0.26) and NR (r = 0.08, p = 0.56) conditions.

To further examine whether the neural measures (i.e., dissimilarity, and connectivity) vary as a function of vividness ratings, we conducted a set of additional analyses for inter-trial neural dissimilarity and network connectivity data for vividness ratings over 8 runs. As shown in **Fig. S1*D*** below, we observed that inter-trial neural pattern distinctiveness in the canonical retrieval-related brain mask in final run was positively correlated with individual’s slope of vividness ratings only for the RP (r = 0.30, p = 0.03) but not NR condition (r = -0.09, p = 0.52). Parallel network-based prediction analysis for average vividness ratings with the same feature selection procedure reported in the main manuscript revealed the prediction values reaching as 0.62 (r_(predicted, observed)_ = 0.62, p < 0.001 by permutation test) from the third run and remaining at a stable level since then. The prediction values outperformed the NR condition as well as the randomly-permutated level (**Fig. S1E**). The selected functional links were superimposed onto a glass brain template for visualization purpose (**Fig. S1F**).

**Associative memory for face-scene associations in the RP, NR and baseline conditions**

To examine the differences in memory performance for remembering the association of face-scene pairs in the RP, NR and baseline conditions, we conducted a 2-by-3 ANOVA for associatve memory of face-scene pairs, with Time (Immediate vs. Delayed) and Condition (RP vs. NR vs. Baseline) as within-subject factors. This analysis revealed a main effect of Condition (F_(2,55)_ = 23.97, *p* < 0.001), and a Condition-by-Time interaction effect (F_(2,55)_= 10.39, *p* < 0.001; **Fig. S2*A***). *Post hoc* t-tests showed significantly better memory performance for RP trials in the delayed than immediate test (*t*_(56)_ = 3.11, *p* = 0.003), but worse memory performance for NR (*t*_(56)_ = -2.53, *p* = 0.014) and baseline items (*t*_(56)_ = -2.45, *p* = 0.018) in the delayed test than the immediate test. These results indicate better memory performance in the delayed than immediate recall test selectively for the RP but not NR condition.

Parallel 2-by-3 repeated ANOVA for false memory scores revealed a main effect of Conditions (F_(2,98)_ = 11.01, *p* < 0.001) (**Fig. S2*B***). Follow-up paired t-tests revealed significantly higher false memory scores in the delayed test for the RP than the NR condition (t_(49)_ = 2.49, p = 0.016), and a marginally significant higher false memory in the immediate test for the RP than NR condition (t_(49)_ = 1.92, p = 0.06). These results indicate that retrieval practice tends to induce higher false memory in both immediate test and delayed test as compared to the NR condition.

**Dynamic changes in neural pattern fidelity over retrieval practice**

To investigate how memory-related neural patterns change over retrieval practice, we computed multi-voxel pattern similarity of neural activity of each run in the canonical retrieval-related brain mask as relative to the initial state of the first run to assess the fidelity of neural patterns in the RP and NR conditions. Paired-t tests revealed significantly higher average neural fidelity across 8 runs in the RP than NR condition (t_(49)_ = 6.99, p < 0.001)(**Fig. S4*B***). Moreover, the higher neural fidelity was positively predictive of better long-term retention gains after consolidation in the RP condition (r = 0.29, p = 0.04; **Fig. S4*C* *left***), but not in the NR condition (r = -0.01, p = 0.95; **Fig. S4*C* *right***). Further statistical analysis revealed a significant difference in these correlation coefficients (Steiger’s z = 1.99, p = 0.023).

We further segmented the overall mask into 15 distinct regions of interest (ROIs) or nodes based on their spatially contiguous voxels, and then examined retrieval-induced changes in neural fidelity over 8 runs in each of these ROIs. As shown **Fig. S5**, we found a general decrease in neural fidelity from the second run to the eighth run (**Fig. S5**). These results indicate that retrieval-induced activity patterns become gradually distinct from the initial state, suggesting a high malleability of memory-related neural representation patterns.

**Dynamic changes in trial-specific neural pattern similarity over retrieval practice**

To investigate how a given memory engram is altered over retrieval practice, we computed the similarity of each trial-specific multi-voxel activity pattern from the second to the eighth run relative to its initial status of the same trial in the first run for the RP and NR conditions separately. As shown in **Fig. S6**A, we conducted a repeated 2-by-7 ANOVA with Run (2^nd^ to 8^th^ run) and Condition (RP vs. NR) as within-subject factors. This analysis revealed a main effect of Condition (F_(1, 49)_ = 33.39, p < 0.001), with significantly higher trial-specific neutral pattern similarity in the RP than NR condition (all t_(49)_ > 2.95, all p < 0.0049). Furthermore, we found a significantly linear decrease in the RP condition (Linear fitting with β = -0.169, p = 0.009; **Fig. S6*B left***), but not in NR condition (β = -0.107, p = 0.098; **Fig. S6*B right***).

Moreover, we conducted correlational analyses to investigate whether retrieval-induced changes in trial-specific neural pattern similarity contribute to subsequent memory outcomes. These analyses revealed no reliable correlations with long-term retention gains (RP: r = -0.19, p = 0.18; NR: r = -0.04, p = 0.78) nor false memory scores (RP: r = 0.17, p = 0.27; NR: r = -0.15, p = 0.34) in RP and NR conditions. The above results indicated that trial-specific neural similarity in the overall brain mask appears higher in the RP than NR condition, but not predictive of subsequent memory outcomes.

**Retrieval-induced dynamic changes in network connectivity in the PPC and MTL**

To further characterize retrieval-induced changes in brain network configurations over retrieval practice, we performed a graph theory-based network analysis for 15-by-15 ROI matrices in the RP (or NR) condition to compute participation coefficients within the MTL, PFC, and PPC for each run. Participation coefficient assesses the diversity of intermodular connections of individual nodes^1^. This analysis revealed a higher participation coefficient for the global network in the last run than the first run (t_(49)_ = 2.20, *p* = 0.03), with most prominent effects for the PPC (t_(49)_ = 2.79, *p* = 0.007) and MTL (t_(49)_ = 2.20, *p* = 0.03) only for the RP condition, but not for PFC (t_(49)_ = -0.14, *p* = 0.89) (**Fig. S7*B***). There was no any reliable network segregation in the NR condition in these three systems (**Fig. S7C**). These results indicate that the PPC and MTL show increased functional segregation over retrieval practice.

**VLPFC connectivity with LPC and other nodes, but not the MTL, predicts long-term retention gains**

To examine the extent to which the VLPFC connectivity with PPC and other nodes is predictive of long-term retentions gains, we first conducted an additional prediction analysis by only taking the right VLPFC functional links with the right LPC into account. The reason of using LPC was because that the LPC node appeared to have relatively high betweenness within PPC nodes (**Fig. 5*D***). This analysis revealed relatively high prediction value in the RP condition (r_(predicted, observed)_ = 0.59, p < 0.001), but not in the NR condition (r_(predicted, observed)_ = 0.08, p = 0.56) (**Fig. S9*A***).

To examine whether the connectivity between VLPFC and PPC is the most crucial predictor, we explored whether the right VLPFC connectivity with the MTL nodes could reliably predict long-term retention gains. This analysis revealed no reliable prediction value for long-term retention gains for the RP (r_(predicted, observed)_ = 0.18, p = 0.22) and NR (r_(predicted, observed)_ = 0.24, p = 0.09) conditions (**Fig. S9*C***). To examine whether the VLPFC connectivity alone is sufficient to predict long-term retention gains, we conducted an additional prediction analysis by only taking the right VLPFC connectivity with other nodes as input features. This analysis revealed a relatively high prediction value of 0.61 (r_(predicted, observed)_ = 0.61, permutated p < 0.001, **Fig. S9*E***), but lower than the one when taking all links into the model (r_(predicted, observed)_ = 0.74, permutated p < 0.001). The selected features were visualized in **Fig. S9*F*** by projecting onto a glass brain template. The right VLPFC links with PPC nodes contribute the most information to predicting long-term retention gains after consolidation. Together, these results indicate that functional connections of the right VLPFC with the LPC_R and other cortical nodes, but not with MTL nodes, over retrieval practice are highly predictive of long-term retention gains.

**Supplementary Figures**

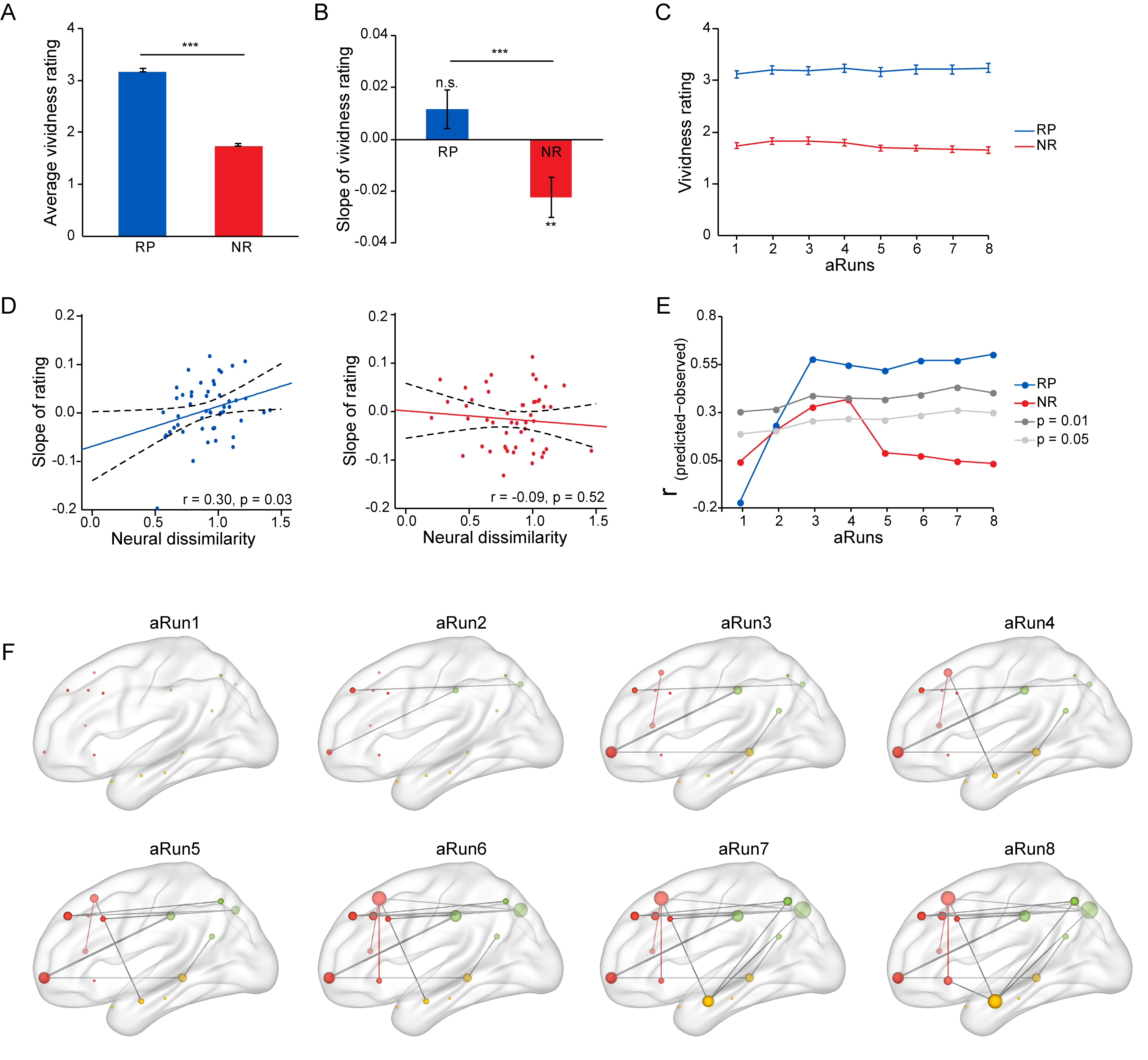

**Fig. S1. Vividness ratings during memory practice over 8 runs and their relations to inter-trial neural distinctiveness and network configurations.** (**A**) Bar graph depicts the average vividness ratings across 8 runs for the RP and NR conditions. (**B**) Bar graph depicts the average slopes of vividness ratings as a function of 8 runs derived from a linear regression model. (**C**) Line plots depict vividness ratings in each run in the RP and NR conditions. (**D**) Scatter plots show the relationship between inter-trial neural distinctiveness in the final run and the slope of vividness ratings for the RP (left) and NR (right) conditions. (**E**) Prediction values for the averaged vividness ratings in the RP (or NR) condition over 8 runs from the top 1% selected links in an accumulative stepwise manner. (**F**) The selected links over 8 runs were projected onto a glass brain template. Notes: ** *P* < 0.01, *** *P* < 0.001, Error bars represent standard error of mean.

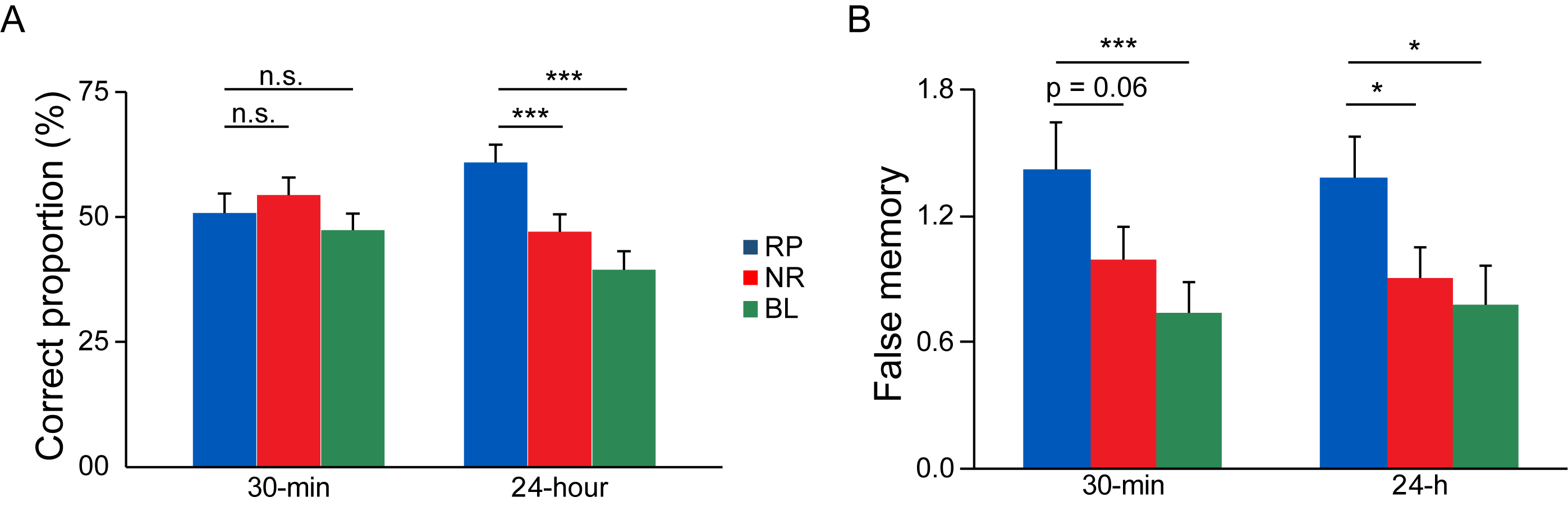

**Fig. S2. Associative memory performance for face-scene pairs in the RP, NR and baseline conditions**. (**A**) Bar graphs depict the average correct proportion for associative memory of face-scene pairs, and (**B**) the average false memory scores – recalled false details of the associated scenes when remembering face-scene associations in the RP, NR and baseline conditions in the immediate (30-minute) and delayed (24-hour) tests. Notes: * P < 0.05; *** P < 0.001; n.s., not significant. Error bars represents standard error of mean.

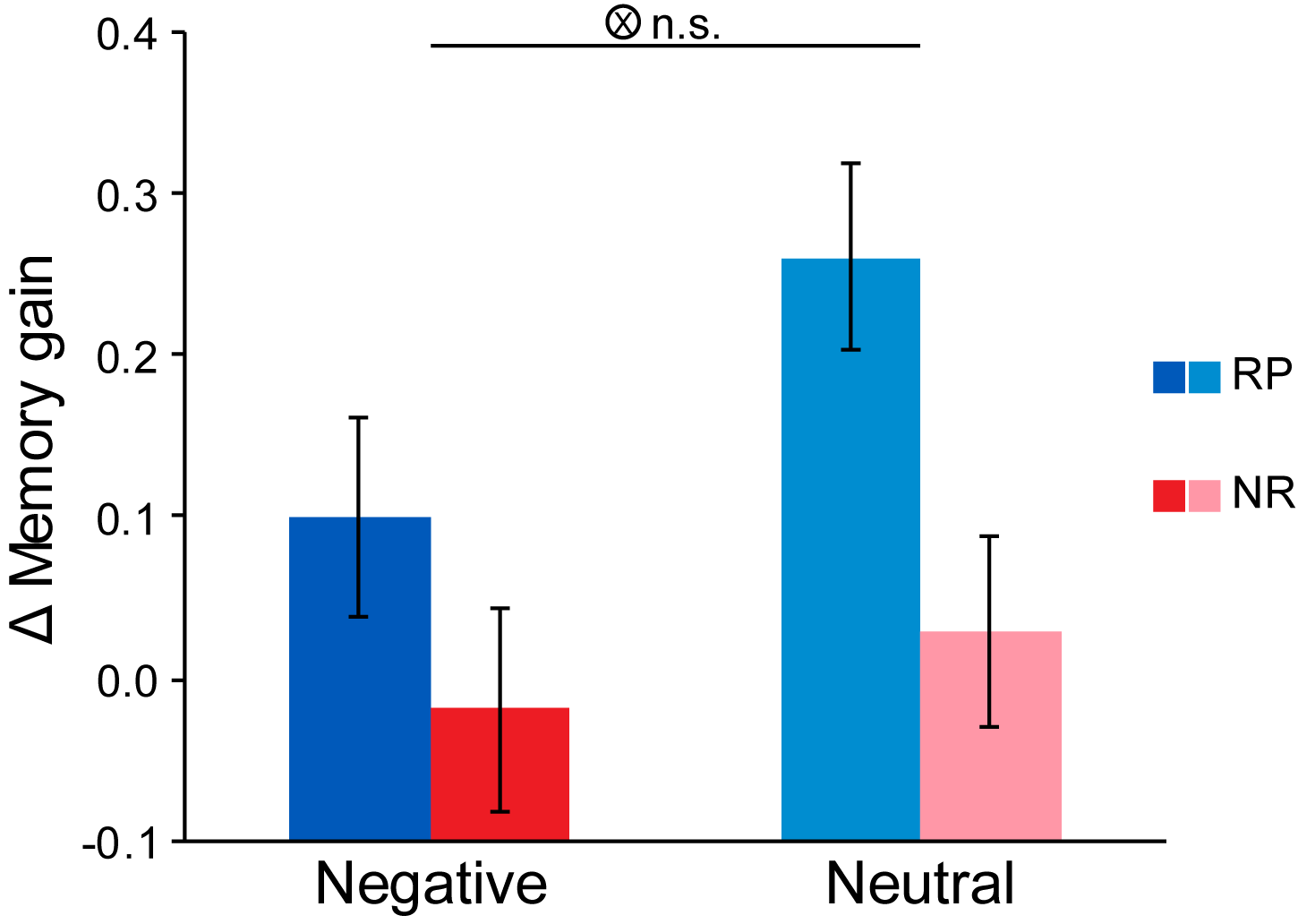

**Fig. S3. Long-term retention gains for face-scene associations with the negative and neutral scenes as stimuli.** A 2-by-2 repeated ANOVA with Valence (Negative vs. Neutral) and Memory (RP vs. NR) as within-subject factors revealed neither the main effect of Valence nor the interaction between Valence and Memory (RP vs. NR) conditions. Notes: n.s., not significant. Error bars represents standard error of mean.

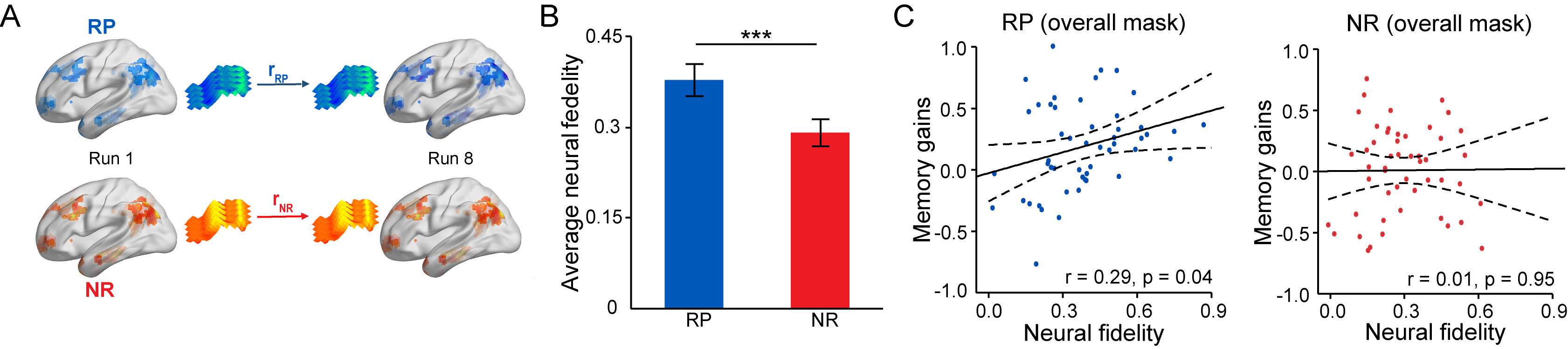

**Fig. S4.** **N****eural pattern fidelity over retrieval practice** (**A**) An illustration of how neural fidelity was computed by Pearson’s correlation in multi-voxel neural activity between the initial and the final state in RP and NR conditions separately. (**B**) Bar graph depicts the average neural pattern fidelity across 8 runs, with significantly higher in the RP than NR condition. (**C**) Scatter plots show a positive correlation of neural pattern fidelity with long-term retention gains in the RP but not NR condition. Notes: *** p < 0.001; Error bars represent standard error of mean.

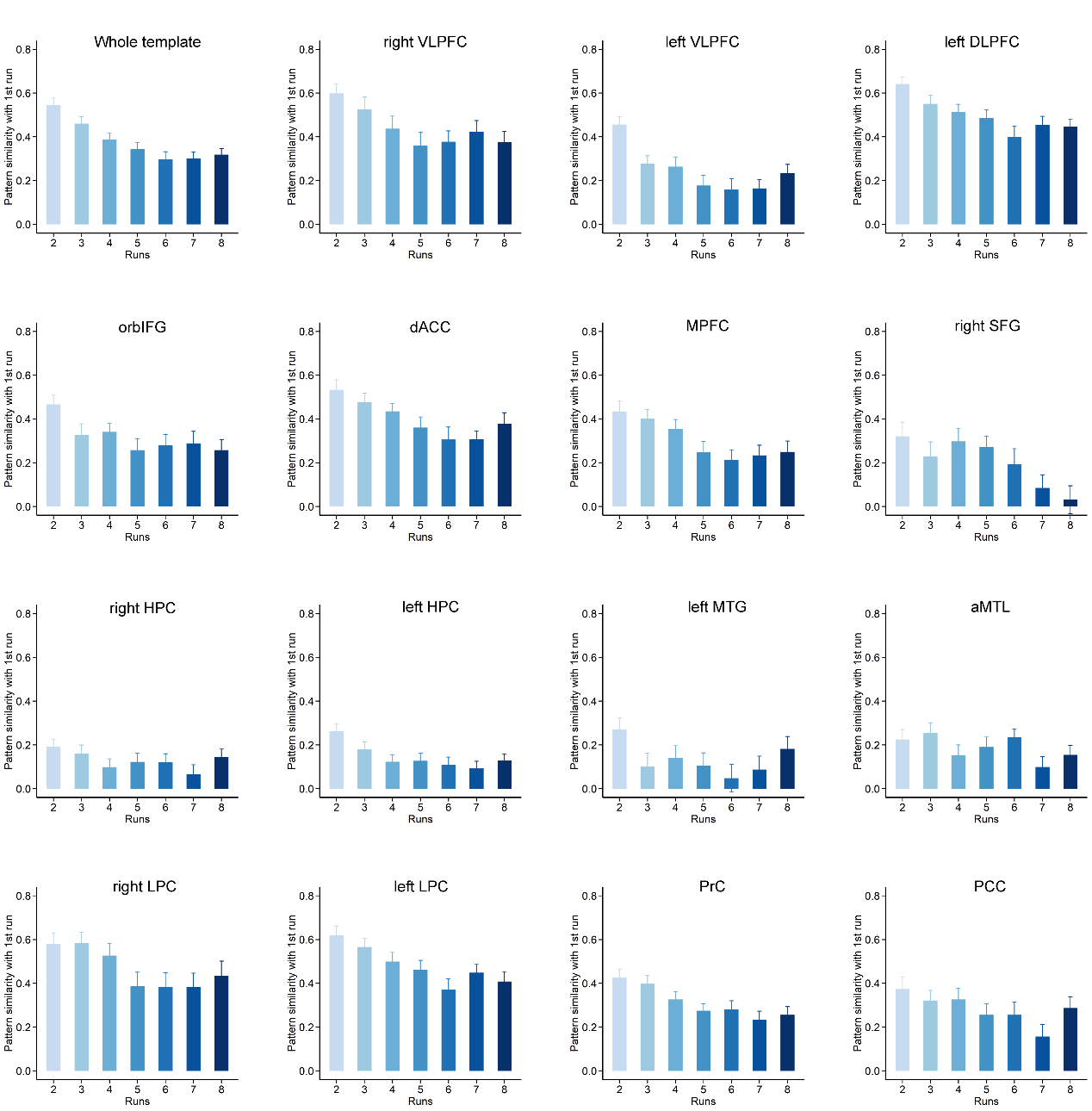

**Fig. S5.** **Multi-voxel neural pattern fidelity over retrieval practice for 15 individual regions.** Bar graphs show a general decrease in neural pattern fidelity from the second run to the eighth run, with most prominent effect in the prefrontal and parietal regions, but relatively weaker pattern in the MTL regions. Notes: Error bars represents standard error of mean.

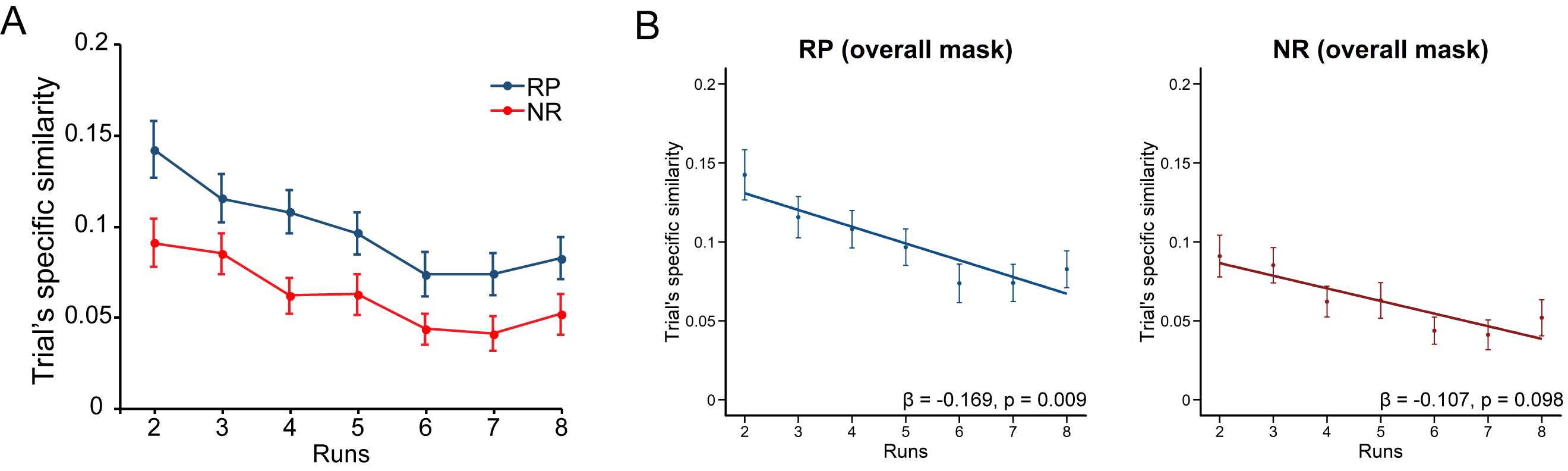

**Fig. S6: Dynamic changes in trial-specific neural pattern similarity over 8 runs.** (**A**) Line charts show changes in trial-specific multi-voxel activity pattern in the canonical retrieval-related brain mask in each run (from 2^nd^ to 8^th^ runs as relative to its initial pattern in the first run for RP and NR conditions separately, with significantly higher in the RP than NR condition. (**B**) Linear regression plots show a significant linear increase in trial-specific neural pattern similarity over 8 runs in the RP but NR condition. Notes: ** p < 0.01. Error bars represent standard error of mean.

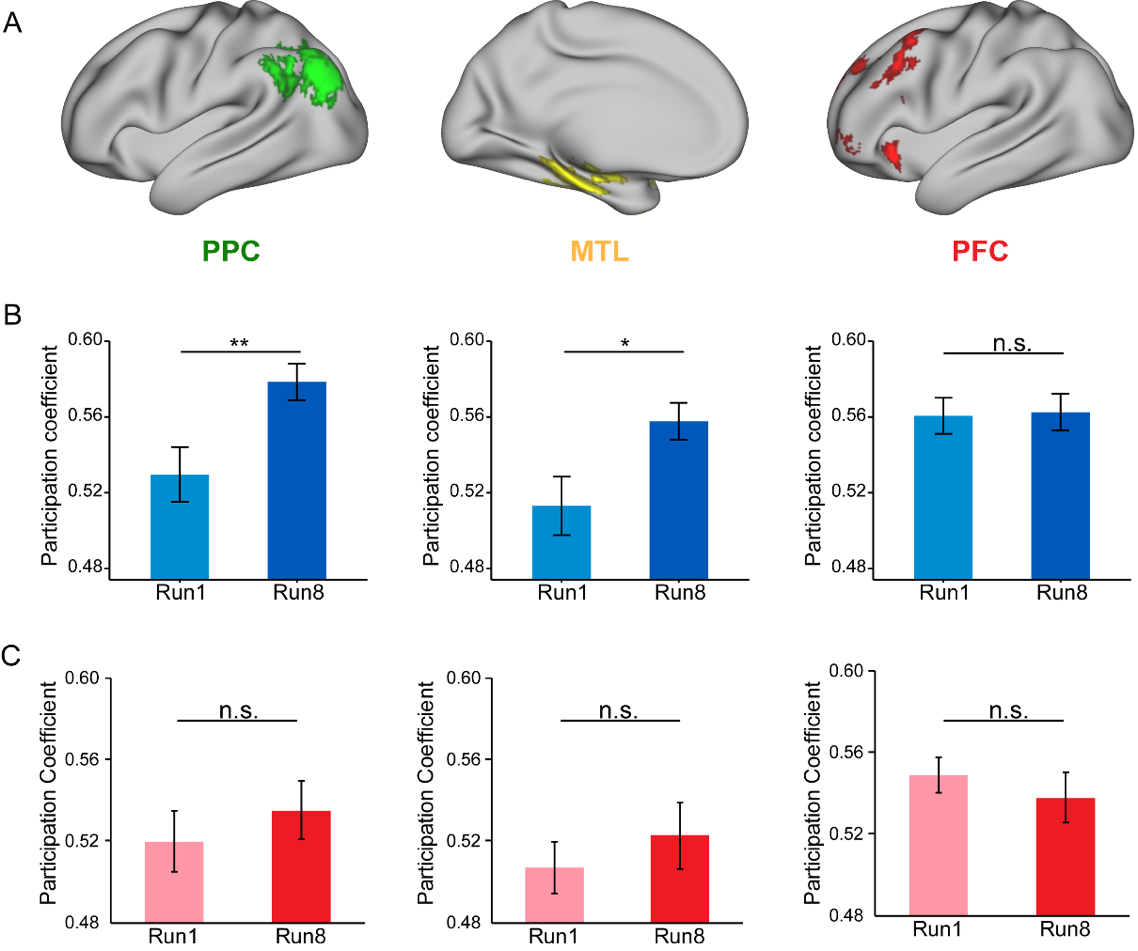

**Fig. S7: Retrieval-induced changes in network configurations in RP and NR conditions. (A)** An illustration of three systems for the PPC (left), MTL (middle) and PFC (right). **(B)** Bar graphs depict participation coefficients between the 1^st^ and 8^th^ run in PPC, MTL, and PFC in RP (B) and NR conditions (C). Notes: n.s., no significant; * p < 0.05; ** p < 0.01.

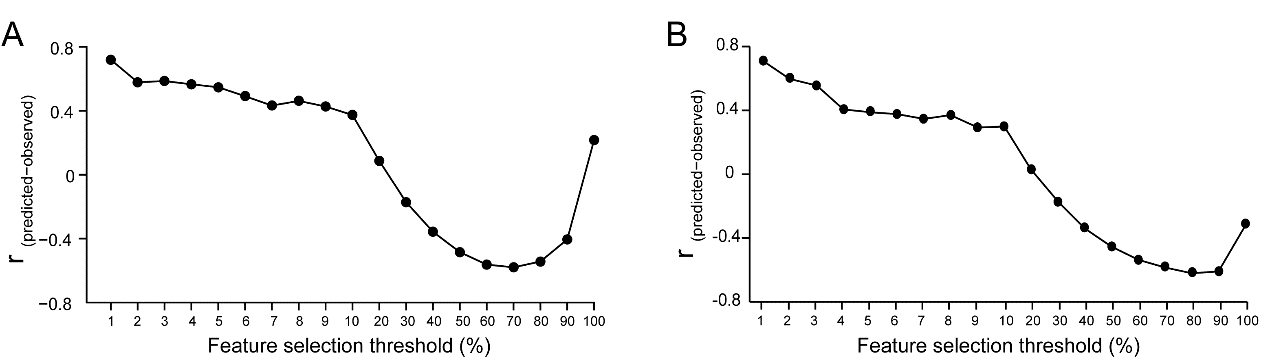

**Fig. S8. Prediction values as a function of different thresholds for feature selection.** Prediction values Information derived from the top 1% links as input features over 8 runs provide the highest accuracy to predict individual’s long-term retention gains after consolidation (**A**) and false memory scores (**B**).

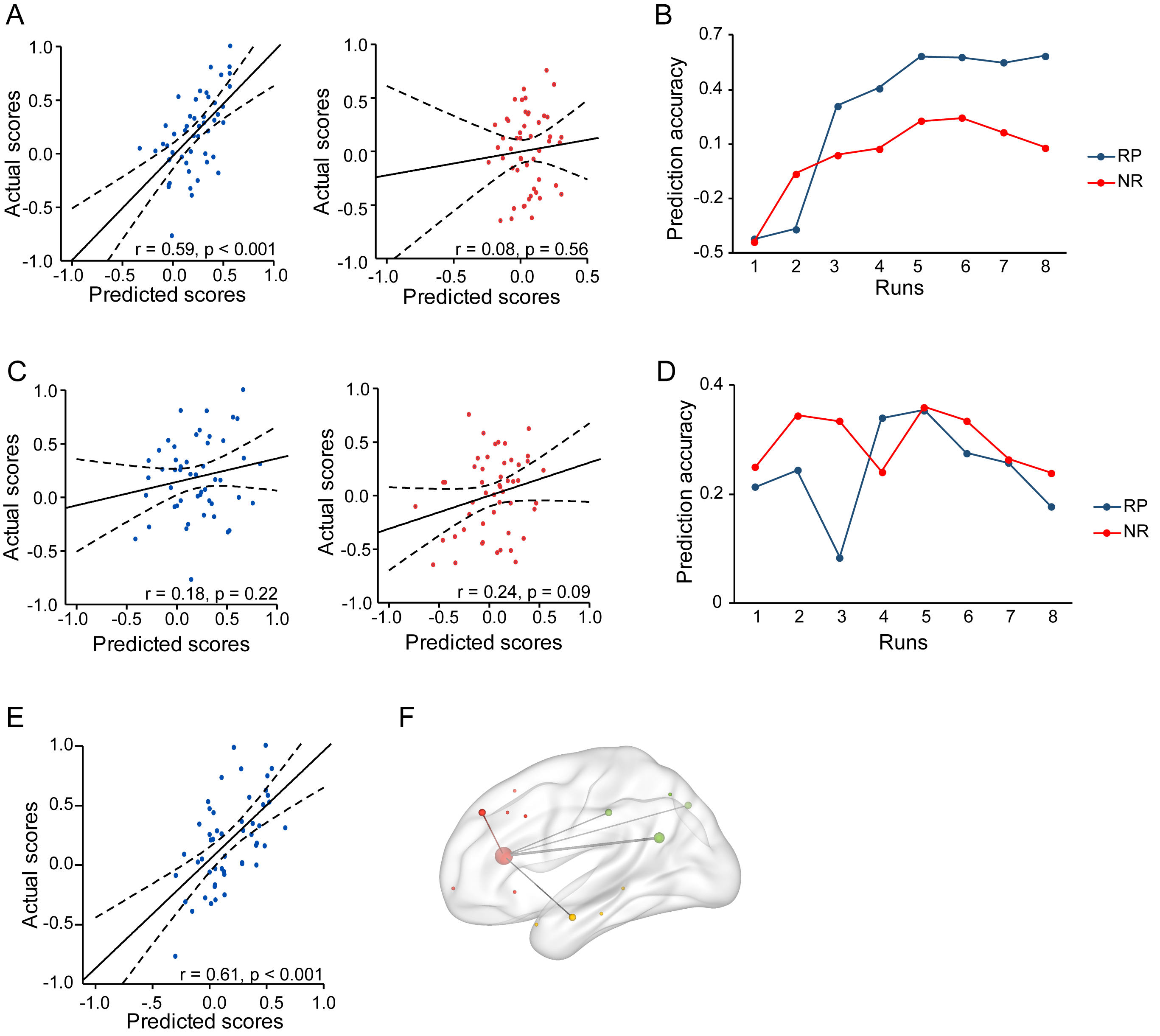

**Fig. S9: Prediction values for long-term retention gains based on the VLPFC connectivity with the LPC, MTL and other nodes.** (**A**) Scatter plots show correlations between observed and predicted scores in RP (left) and NR (right) conditions and (**B**) line charts show prediction values as a function of 8 runs in the RP condition which outperformed the NR condition, based on the right VLPFC functional connections with the right LPC as input features. (**C**) Scatter plots show correlations between observed and predicted scores in RP (left) and NR (right) conditions, and line charts show prediction values as a function of 8 runs in the RP condition which did not differ from the NR condition, based on the VLPFC functional connections with the MTL nodes as input features. (**D**) line charts show prediction values as a function of 8 runs in the RP condition which outperformed the NR condition, based on the right VLPFC functional connections with the right MTL as input features. (**E**) Scatter plots show correlations between observed and predicted scores based on the VLPFC connections with other nodes as input features, (**F**) the selected links were visualized by projecting onto a glass brain template.
